# Iterative sacrificial 3D printing and polymer casting to create complex vascular grafts and multi-compartment bioartificial organs

**DOI:** 10.1101/2024.09.29.615298

**Authors:** Jonathan A. Brassard, Saleth Sidharthan Dharmaraj, Hamid Ebrahimi Orimi, Daria Vdovenko, Yannick Rioux, Hanwen Wang, Kurtis Champion, Florent Lemaire, Brenden N. Moeun, Jia Zhao, Shenghui Liang, Timothy J. Kieffer, Jean I. Tchervenkov, Gilles Soulez, André Bégin-Drolet, Jean Ruel, Richard L. Leask, Steven Paraskevas, Corinne A. Hoesli

## Abstract

Several emerging strategies to engineer artificial organs employ 3D printing to create vascular templates to provide nutrients and oxygen to immobilized cells. Significant challenges emerge when considering clinical implementation such as immune rejection of allogeneic cell sources, as well as achieving adequate perfusion and integration with endogenous vasculature. We propose a method by which cell-laden hydrogels are molded around ready-made polymeric vascular templates created via 3D printing to create human-scale artificial organs with internal vasculature. We applied this technique to create bioartificial pancreas systems with up to 9 internal flow channels via sacrificial carbohydrate glass 3D printing, porogen-loaded polycarbonate polyurethane dip-coating, followed by casting cell-laden hydrogels around the vascular templates. We optimized porogen size and concentration to maximise the porosity of our scaffolds without compromising mechanical properties, resulting in suture retention strength and compliance respectively matching commercial vascular grafts and native vessels. Bioreactor perfusion studies showed survival and maturation of stem cell derived pancreatic islets without significant differences to traditional suspension culture protocols. Insulin response dynamics were rapid in response to a glucose challenge at the perfusion inlet. Transplantation of the devices as iliac arteriovenous shunts in nondiabetic pigs confirmed safety and patency. These results show promise for the development of an implantable vascularized pancreas for the treatment of type 1 diabetes and demonstrate how bioartificial organs with engineered vascular geometries can be designed for translational applications.

## INTRODUCTION

Cellular therapy is revolutionizing how we treat degenerative disease. While most cellular therapies approved by regulatory agencies rely on blood-derived products, many pluripotent stem cell-derived therapies to replace failing tissues and organs are in the pipeline^1^. The effective delivery of many of these products requires organization into artificial tissues that can sustain long-term graft function. At tissue densities, the maximum distance from a blood vessel to a transplanted cell to ensure adequate oxygenation is typically on the order of 0.1∼0.5 mm. To solve this molecular transport problem, physiological systems have evolved to comprise dense hierarchical vascular architectures. Artificial organ systems integrating vascular lattices have been engineered using layer-by-layer 3D printing, cell casting around sacrificial self-supporting vascular templates^2^, and *in situ* creation of perfusable networks either through embedded writing^3–5^ or photochemical reactions^6^. However, clinical translation of these strategies will require careful consideration of tissue scale and geometry, as well as implantation and physiological integration strategies.

As an example, pancreatic islet transplantation, an established cellular therapy to treat type 1 diabetes, currently relies on intraportal infusion of ∼8,000 islet equivalents per kg body weight (∼900 million cells for a 75 kg recipient) to achieve blood glucose normalization for at least one year in most recipients. The therapy is currently limited by ∼100-fold insufficient cadaveric islet supply, as well as the need to maintain lifelong immunosuppression to limit allograft rejection^7^. Immune suppression regimens applied increase risks of infection, increase the incidence of cancer, and can also be harmful to islet survival and function. With pluripotent stem cell-derived islets emerging as virtually unlimited cell source, there has been renewed interest in engineering extra-hepatic transplantation sites. Macroencapsulation devices are particularly promising for stem cell-derived islet delivery due to their ease of monitoring and their potential to be retrieved. However, because these devices rely on passive diffusion and the transport of oxygen and nutrients is impeded by the physical barrier protecting the highly metabolic islets, the geometry of the devices is restricted to a flattened pouch containing a limited cell mass to avoid excessive cell death and loss of function^8^. Scaling to high cell density and human therapeutic doses has remained a significant challenge using these traditional approaches, with surface areas estimated on the order of 1 m^2^ needed for the survival of a therapeutic dose of islets at 30 mmHg of oxygen^9^.

So far, diffusion-based bioartificial pancreas devices such as microbeads, porous polymer pouches, or diffusion devices with an oxygen compartment, all demonstrated safety but limited efficacy in clinical trials^10,11^. The foreign body response to the implanted materials, the time required to establish neovasculature in humans, as well as insufficient vascular densities may all contribute to limited therapeutic impact^12^. To facilitate oxygen delivery and provide a better microenvironment for the transplanted islets, several strategies have aimed at enhancing vascularization, either via a pre-vascularized bed^13^ or by enhancing graft integration via capillary ingrowth^10,14^. Unfortunately, these vascularization strategies are incompatible with physical immunoprotection, making vascularization superficial and inefficient when used in combination with immunoprotective macroencapsulation devices.

To address the challenges associated with passive diffusion, some approaches have used a refillable oxygen chamber^15^, oxygen generating materials^16^, an external pump for convective transfer^17^ or ultrafiltration^18,19^. Oxygenation supply can significantly improve long-term islet survival, but the amount of insulin reaching the circulation may not have significant therapeutic impact if vascular perfusion is insufficient^11^. Alternatively, direct anastomosis to the arterial system of the host using a vascular graft surrounded by encapsulated islets has been proposed either with^20^ or without^21^ an immunoisolating component. The strategy developed by W.L. Grace was effective in normalizing blood glucose in dogs for up to a year when two devices with allogeneic islets were transplanted per animal, but failure at the device-vascular graft interface led to sudden death in preclinical studies, putting this strategy largely on hold since the 1990s^20,22,23^. New strategies that can eliminate the many material interfaces involved in device fabrication and avoid the need to transplant multiple devices could improve the safety and efficacy profile of this approach.

We propose a novel encapsulation strategy whereby therapeutic cells in a hydrogel support are introduced in a ready-made pouch with internal vascular polymeric networks that can be directly anastomosed to the recipient circulation. The pouch and internal vascular lattice geometries are created by 3D printing or casted sacrificial templates to achieve customizable vascular network device designs required to scale up to human therapeutic doses. We hypothesized that such a device would not only eliminate the need for host vascular ingrowth, but also enhance cell survival and insulin release efficiency by harnessing the convective power of circulating blood flow. In addition, our design process enables the generation of a seamless device made from only polyurethane that can be sutured directly to blood vessels with good compliance matching, suture retention strength and adaptability.

We first optimized the internal vascular network by maximizing diffusion through the channel walls while maintaining mechanical properties that would be suitable for suturing and native blood pressure. We demonstrate biocompatibility in mice with limited fibrosis and allowing for the formation of an endothelium in the inner surface. Using an *in vitro* vascular chamber, we show that beta cell insulin secretion responses are rapidly detected at the device outlet after increasing glucose concentration at the inlet. We also characterized the oxygen profile within our device and evaluate the performance of the device depending on the magnitude of convective mass transport. Moreover, stem cell derived islets are shown to survive and mature inside the perfused device. Finally, arteriovenous shunt implants in hybrid pigs confirmed that the devices are safe and patent up to 1 week after implantation.

## RESULTS

### Fabrication of highly porous vascular grafts

The intended devices comprise a vascular interface and a cell immobilisation compartment which should meet requirements both as a vascular graft and cell encapsulation system. Ideally, the graft material mechanical properties such as compliance, tensile strength and burst pressure would match native vessels, and its surface properties would avoid thrombosis and an inflammatory response. When considering cell immobilisation features, the design should allow delivery of a therapeutic cell dose in a compact geometry. To address this, we envisioned a fabrication process that would create complex internal vascular networks with high mass transport capacity through its wall. To avoid cell death during assembly, the device should be ready-made and sterilized, allowing cell loading shortly before implantation. The device should allow filling, and rapid *in situ* gelation of a cell immobilisation material, such as a hydrogel, supporting cell survival and function. The cell immobilisation material should integrate features such as acting as a physical barrier against immune cell infiltration, immunomodulation, or integration of moieties that favour graft survival or function.

With these constraints in mind, we designed a pipeline where sacrificial carbohydrate is first used to create a vascular network template^2,24,25^ that can be dip coated in a polymeric solution to generate complex vascular grafts (Fig. 1a and Supp. Figure 1). To maximize the transport of nutrient and oxygen through the vascular interface, we included sodium chloride in the solution to act as a porogen, leaving interconnected pores inside the graft material after dissolution. We used a commercial polycarbonate polyurethane (Chronoflex AR) as the graft material because it is biocompatible, has good elongation under stress and excellent tensile strength, and can be dissolved in organic solvents^26,27^, making it an ideal material for making hierarchical graft via dip coating.

**Fig 1.**
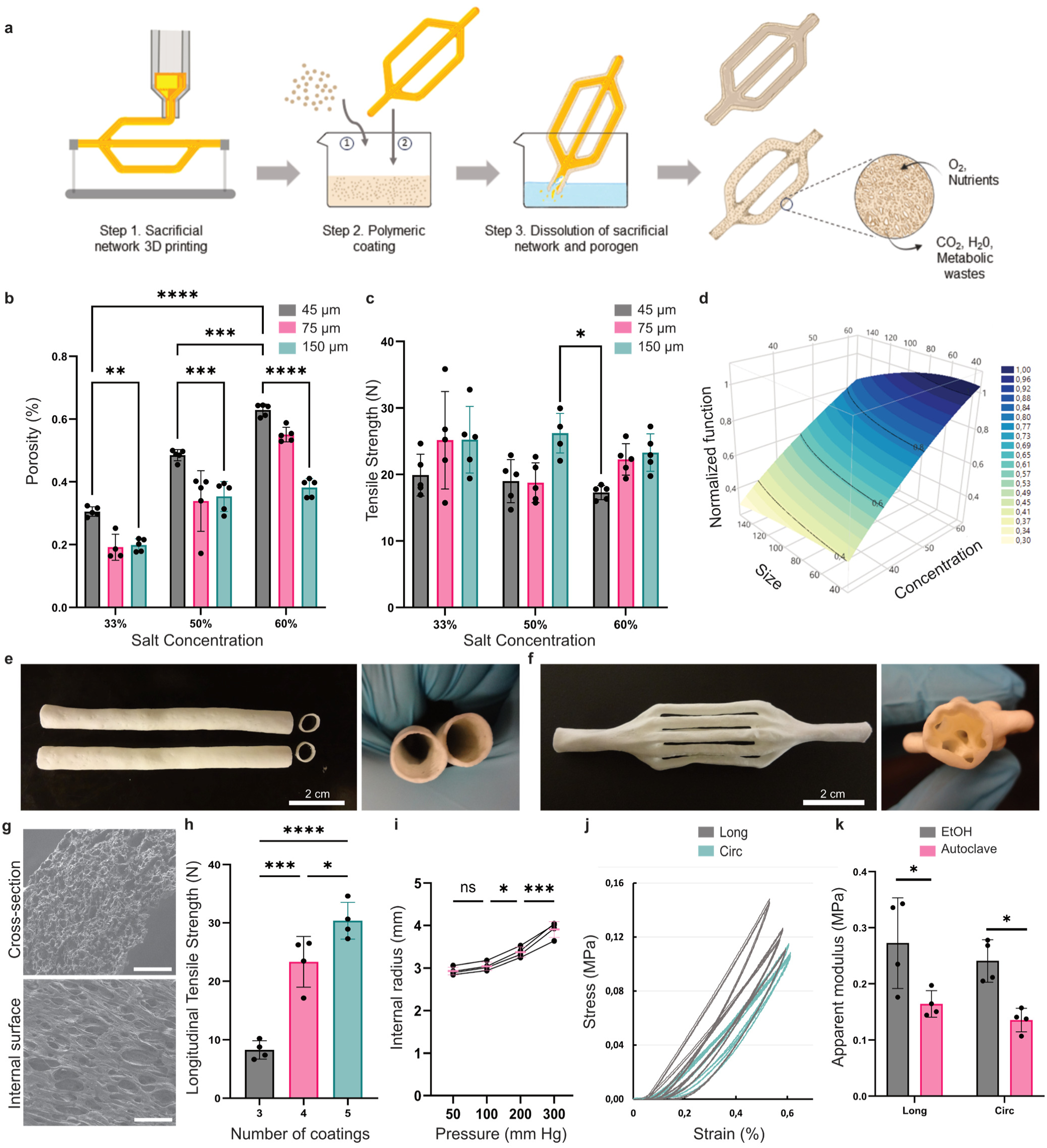
Characterization of vascular grafts. **a**, Illustration of the graft fabrication process. **b**, Graph showing porosity of the grafts based on salt size and concentration. Mean and standard deviation of five different replicates are shown for each condition. **p < 0.01, ***p<0.001, ****p<0.0001, determined by two-way ANOVA with Tukey’s multiple comparisons test. Additional p-values for Tukey’s test are in Supplementary Table 1. **c**, Graph showing longitudinal tensile strength of the grafts based on salt size and concentration. Mean and standard deviation of five different replicates are shown for each condition. *p<0.05, determined by two-way ANOVA with Tukey’s multiple comparisons test. **d**, Normalized surface response for the optimization between the porosity and the tensile strength of the grafts. **e-f**, Representative images of the single-channel (e) and 9-channel (f) grafts after dissolution of the sugar network and porogen. **g**, Representative scanning electron microcopy images of the cross-section and internal surface of the grafts. Scale bars, 200 µm. **h**, Graph showing the increase of longitudinal tensile strength in function of the number of layers. Mean and standard deviation of four different prosthesis are shown. ****p < 0.0001, ***p<0.001, *p<0.05, determined by one-way ANOVA with Tukey’s multiple comparisons test. **i**, Graph showing the increase of internal radius of graft depending on the pressure. Mean and standard deviation of four different grafts are shown. ***p<0.001, *p<0.05, determined by one-way ANOVA with Tukey’s multiple comparisons test. **j**, Stress-strain curves obtained using a biaxial tensile tester, showing 3 successive extension and release for 4 different prostheses. Circumferential and longitudinal axis are compared. **k**, Graph showing apparent modulus in function of the axis (longitudinal vs circumferential) and the method of sterilization (Ethanol or autoclaving). Mean and standard deviation of four different grafts are shown. *p<0.05, determined by two-way ANOVA with Tukey’s multiple comparisons test.

We applied a 3 x 3 full factorial design to determine the effect of porogen particle size and concentration on the properties of the dip coated prosthesis, including reproducibility of the coating, dried material porosity, swelling of the graft, and ultimate longitudinal tensile strength. The volumetric porosity of the prosthesis increased with increasing salt concentration and decreasing salt size (Fig. 1b, Supp. Figure 2a-b), while the tensile strength followed an opposite trend (Fig. 1c, Supp. Figure 2c-d). Surprisingly, the tensile strength of the graft was not significantly affected by the porogen concentration (p=0.14), potentially reflecting a more homogeneous coating at smaller porogen size and higher porogen concentration due to higher viscosity during the dip coating process. Indeed, more polymer was deposited on the sugar template at higher porogen particle diameters or concentration (Supp. Figure 2e). Due to this, the desirability function was weighted more strongly in favor of porosity compared to tensile strength (70% vs 30%) in anticipation of nutrient and insulin transport requirements. The graft conditions (45 µm size & 60% concentration; Fig. 1d) were chosen based on this analysis and used for the rest of the experiments. Because of extensive swelling of the grafts (Supp. Figure 2f), the diameter of sugar constructs for the following experiments was lowered to 5 mm to achieve target internal diameters of 6 mm at the vascular graft inlet and outlet for anastomosis to pig or human circulation.

With the selected dip coating conditions, we then validated that the polymeric solution could allow robust coating of complex hierarchical sugar networks. We successfully fabricated single tubular grafts as well as grafts with 9 branches that can be simultaneously irrigated (Figure. 1e-f). The porosity of the graft was 63 ± 2% in dry conditions (Figure 1b), and pores were visible throughout the material as well as at the surface (Figure. 1g). During characterization, we found that three coating layers produced grafts that had defects leading to early rupture, whereas four or five coatings produced strong prostheses with a longitudinal tensile strength above 20 N (Figure. 1h). We then quantified the radial expansion of the graft under increasing pressure (Figure. 1i), resulting in compliance values in the range of native vessels: 7.47 ± 0.64 %/100 mmHg at a 50-100 mmHg range ^28^. Using a biaxial tensile tester, we show that the grafts have viscoelastic characteristics know for blood vessels in both the longitudinal and circumferential direction (Fig. 1j). Although autoclaving the graft decreased its apparent modulus, the material kept sufficient strength for intervascular implantation (Fig. 1k).

The graft material exhibited similar platelet adhesion when compared to commercially available Teflon graft (Supp. Figure 3c-d). Subcutaneous implantation of cross-section of the graft in immunocompromised mice did not trigger a strong fibrotic response, as shown by limited fibrous capsule seen by collagen staining with Sirius Red (Supp. Figure 3e-f). After 28 days, substantial vascular structures could be seen inside the pores of the graft, suggesting that some degree of vascularization inside the graft could also enhance the delivery of oxygen and nutrients to the encapsulated cells in the final device. If needed, additional hemocompatibility could be provided by luminal endothelialisation either through endothelial cell pre-seeding or post-implantation endothelial colony forming cell capture^29^. Preliminary studies indicate that a homogeneous endothelium can be established on graft luminal surfaces (Supp. Figure 3a-b).

### Vascularized macroencapsulation device fabrication

After establishing the method to create the vascular template, an external compartment was introduced via an analogous sacrificial material casting and polymer dip-coating strategy (Fig. 2a, Supp. Figure 2b). One important requirement of the technique is the independence of the sacrificial material, the porogen and the coating polymer. In this case, polycarbonate polyurethane can be dissolved in organic solvents (DMF and DMAC) that minimally interact with the carbohydrate glass and porogen used as a sacrificial material. To avoid a build-up of stress created by uneven deformation of the material, the pouch surrounding the prosthesis was created using the same combination of porogen and polymer, resulting in a seamless encapsulation device with an internal perfusable network (Fig. 2b-c). Both the vascular graft and device are highly permeable to fluorescently labelled dextran molecules (20kDa) when porogen is incorporated in the fabrication process (Figure 2d, Supp. Movie 1), suggesting that the vascular interface material would not significantly delay oxygen or insulin transport. We then confirmed that devices would sustain the mechanical forces exerted from suturing and blood flow. Although elongation at rupture was slightly decreased, the longitudinal tensile strength of the devices was increased, ensuring both the elasticity and strength expected of a vascular graft replacement (Fig. 2e). Suture retention strength was comparable to commercially available graft (4.2 ± 0.4 N vs 3.6 ± 0.5 N), showing promise for device implantation (Fig. 2f). The resulting devices possess burst pressure that can withstand blood pressure in the artery, specifically when they are filled with hydrogel (Fig. 2g)^30^.

**Fig 2.**
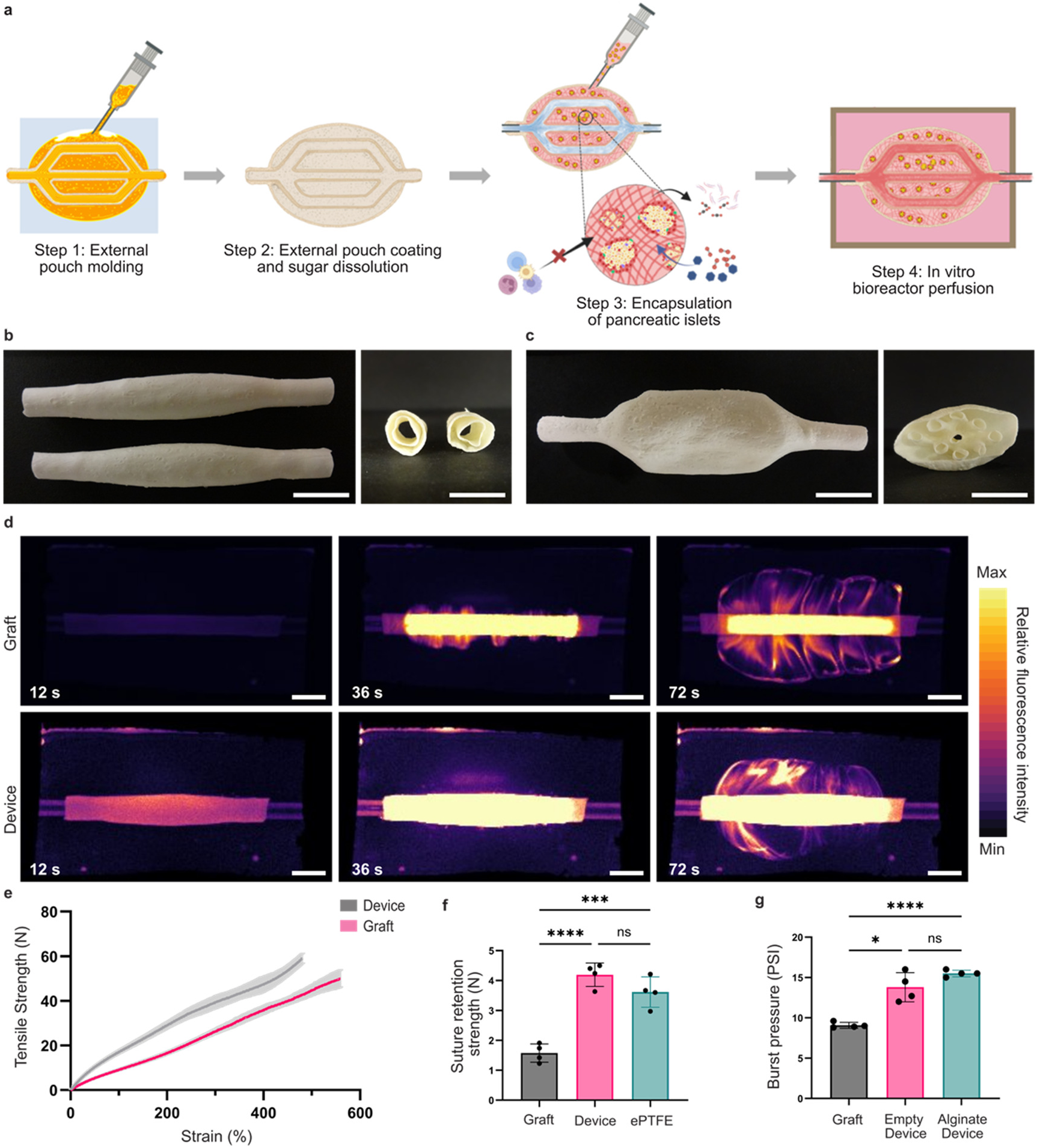
Characterization of device fabricated by iterative 3D printing and polymer casting. **a**, Illustration of the fabrication process of the device and subsequent culture in bioreactor. **b-c**, Photographs showing single channel (b) and 9-channel (c) devices and their cross-section. Scale bars, 1 cm. **d**, Fluorescent images taken using a small animal imager showing the diffusion of fluorescently labelled 20 kDa dextran through a graft alone (top) or an empty device (bottom). Scale bars, 1 cm. **e**, Graph showing force-strain curves comparing the graft alone (pink) vs empty single channel device (grey). Standard deviation of 3 independent replicates is shown in light grey. **f**, Graph comparing the suture retention strength of the graft and empty device to commercial ePTFE alternative. Mean and standard deviation for four different samples are shown. ****p < 0.0001, ***p<0.001, determined by one-way ANOVA with Tukey’s multiple comparisons test. **g**, Graph showing the burst pressure of the prosthesis alone or the device with and without alginate filling. Mean and standard deviation of four different samples are shown for each condition. ****p < 0.0001, *p<0.05, determined by one-way ANOVA with Dunn’s multiple comparisons test.

For their use *in vivo*, device pouches can be filled with cells and various hydrogel precursors, either with the goal of promoting *in situ* vascularization or to confer immunoprotection. Alginate is an anionic polysaccharide that can form a hydrogel when supplied with cationic ions. Even at high polymer concentrations, alginate gelation is cell friendly and does not elicit a significant immune response, explaining its popularity as a material for islet encapsulation^8^. Alginate has thus been used to protect islet in many different settings, including reversal of hypoglycemia by allografts in immunocompetent mice^31,32^ and in non-human primates^33^. Using our modular device approach, cells can be mixed with alginate precursors and injected inside the device prior to gelation. The 5% alginate (50:50 LVM:MVG mixture) composition selected was previously shown to have low antibody permeability, low fibrotic overgrowth, and be stable for up to 5 months in rodent models^32^. After hydrogel formation, alginate could serve as an immunoisolating component while the internal network and external pouch will respectively serve as nutrients sources and device support. To validate the survival of the cells, we designed a bioreactor that can host the device under perfusion when connected to a peristaltic pump (Fig. 3a-b, Supp. Figure 4). Encapsulated MIN6 cell aggregates survived after several days of perfused culture (Fig. 3c). Step changes in glucose concentration triggered rapid release of insulin from the immobilised cells (Figure 3d), confirming the physiological relevance of our approach in minimizing mass transport delay using convective transport. We then tested large-scale devices containing 9 channels to allow increased mass transport through the device (Fig. 3e). Perfusion for 48-hours showed that cells near the channel (<500 µm) survive, creating a cell survival profile that reached a substantial portion of the device cross-section (Fig. 3f-h).

**Fig 3.**
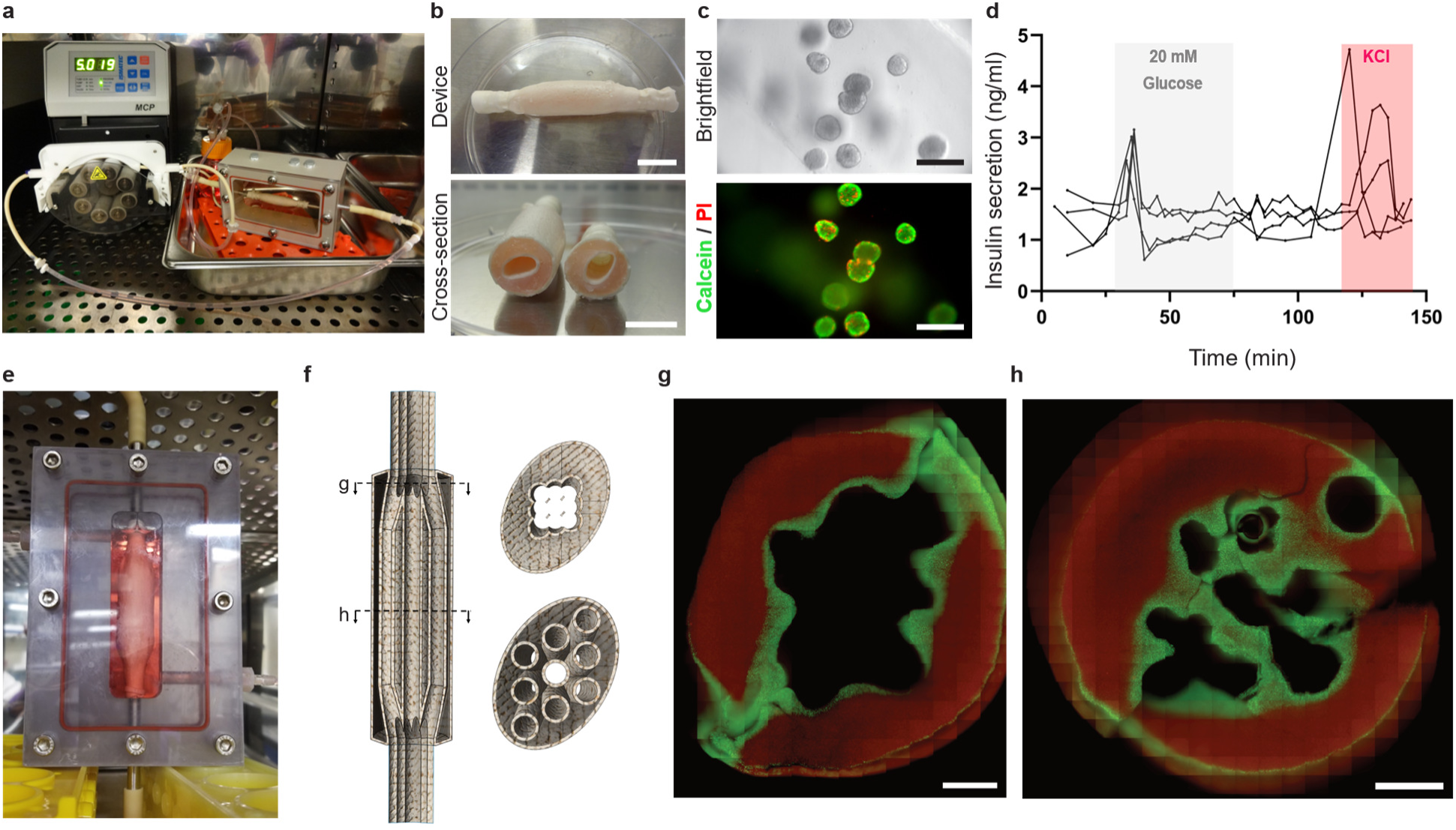
Assessment of survival and function during *in vitro* perfusion. **a**, Perfusion set-up for culture inside our custom-made bioreactor. **b**, Retrieval of the device after 48 hours of perfusion showing unobstructed lumen with stable alginate hydrogel inside the external compartment. Scale bars, 1 cm. **c**, Representative live/dead staining of perfused MIN6 aggregates with Calcein AM (green) and propidium iodide (red). Scale bars, 200 µm. **d**, Insulin secretion in response to step change in glucose concentration or KCl addition for devices with MIN6 cells. The graph shows the insulin release curves for four independent experiments. **e**, Macroscopic image of the perfusion of a 9-channel device. **f**, Illustration for section of the device imaged in (g) ang (h). **g-h**, Representative live/dead staining of perfused MIN6 aggregates in the 9-channel device, with Calcein AM (green) and propidium iodide (red). Scale bars, 4 mm.

### Modelling of oxygen consumption

To inform the design of our current and future devices, a computational model was developed to predict the oxygen profiles through the flow channels and the cell loaded compartment. A single channel device was modelled using the physical characteristics measured from our first iteration device, and the model parameters were taken from the literature and previous studies from our laboratory (Supplementary Table 4). As one of the main limiting components for cellular viability in macroencapsulation devices, mass transport of oxygen through the graft wall and alginate hydrogel was calculated for different flow rate and varying cell density. We previously reported that MIN6 cells show a sharp decline in cell viability at oxygen partial pressure (pO_2_) values of ∼3 mmHg^34^. As expected, hypoxic zones progressively widen with increasing cell densities, reducing the maximum thickness of the device that would prevent cell apoptosis (Fig. 4a-c). Our experimental setup yields a viable cell zone of 522 ± 64 µm for 20 million MIN6 cell/mL, versus 361 ± 77 µm for 40 million MIN6 cell/mL, values that agree with the theoretical model (Fig. 4d-e). In addition to consumption of oxygen radially, the simulated laminar flow through the device could be expected to show reduced availability of oxygen at the outlet compared to the inlet due to the effect of both convective transport and device length on oxygen distribution. To further investigate the importance of convection on the oxygen profile through the devices, we performed oxygen measurements at the outlet of the device at different flow rates (Fig. 4f), while maintaining the oxygen concentration at the inlet constant (Ambiant pO_2_). Below 25 mL/min, we show a progressive decline in oxygen concentration at the outlet that matches the decline seen in our theoretical model (Fig. 4g). Above that threshold, oxygen consumption compared to the flow rate was negligeable (outlet/inlet concentration ratio ≈ 1), confirming that an arteriovenous shunt in large animal and human would provide enough flow to prevent substantial drop in oxygen concentration through the device. These results highlight not only the added value of leveraging convection for mass transfer in encapsulation devices, but also the design constraints for the creation of more complex macroencapsulation devices with multiple branches in the internal network.

**Fig 4.**
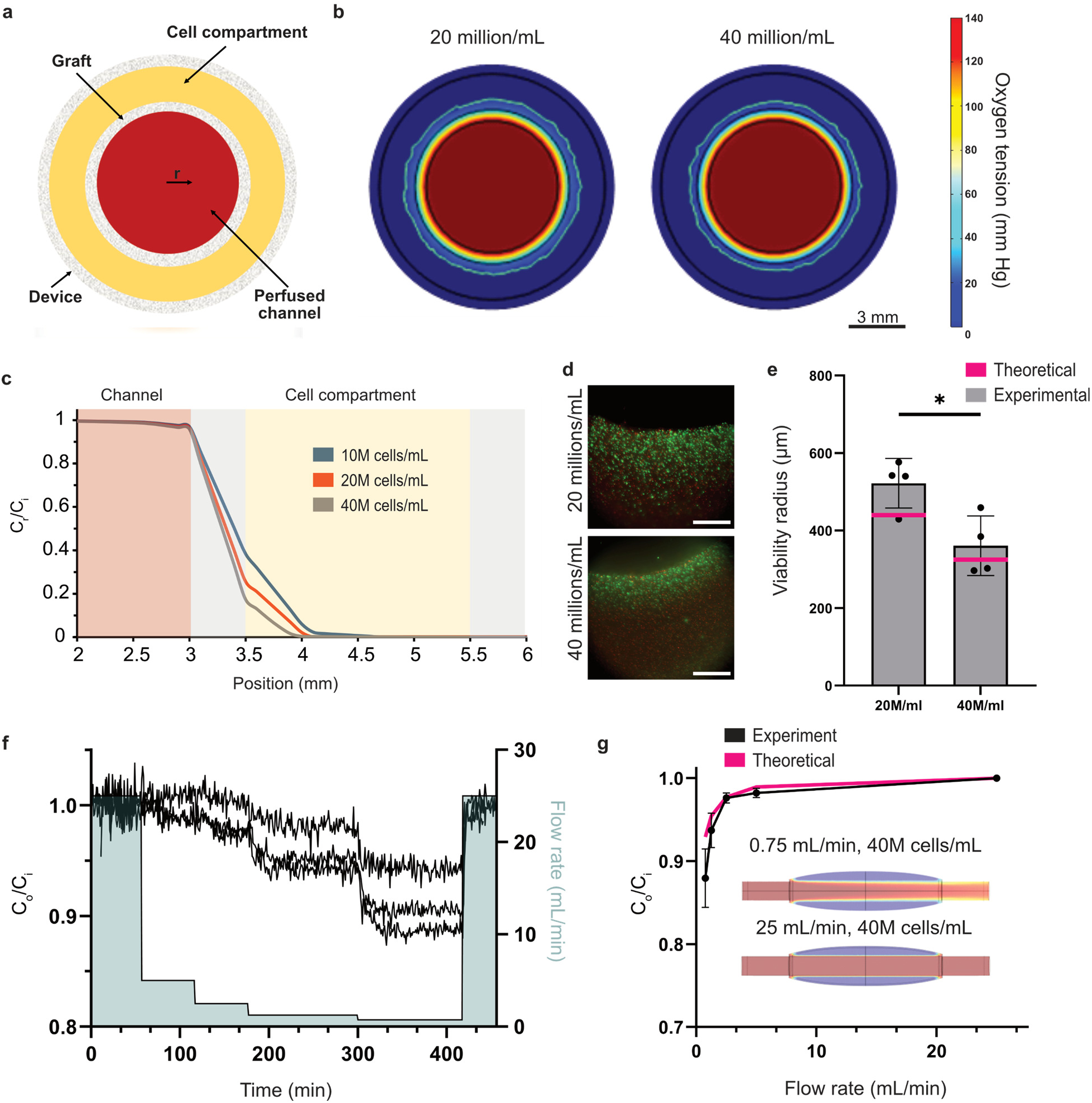
Oxygen profiles through device filled with MIN6. **a**, Schematic representation of a single-channel device cross section. **b**, Corresponding oxygen partial pressure profiles predicted by a finite element model. The white contour represents the limit for necrotic oxygen tension (0.1 mm Hg). **c**, Finite element model predictions of oxygen partial pressure radial profiles at different MIN6 cell concentrations. **d**, Representative live/dead staining of MIN6 cells with Calcein AM (green) and propidium iodide (red). Images were obtained 48 hours after seeding MIN6 in devices perfused at 25 mL/min. **e**, Distance from the internal channel wall at which MIN6 cells transition from high to low viability. Mean and standard deviation for four independent experiments and value for the theoretical model (pink) are shown. *p<0.05, determined by an unpaired t-test. **f**, Experimental data showing the effect of flow rate on the oxygen concentration at the outlet of the device. Perfusion of a device with 40 million cells/mL. **g**, Effect of flow rate on the theoretical and experimental oxygen concentration at the outlet of the device at 40 million cells/mL loading. Illustration inside the graph shows longitudinal oxygen profiles in function of flow rate predicted by a finite element model. C_i_ = oxygen concentration at the inlet, C_r_ = oxygen concentration at different radius, C_o_ = oxygen concentration at the outlet.

### Stem cell derived islet survive and mature during *in vitro* perfusion

Stem cell derived islets (SC-islets) are a promising insulin producing cell source for cell therapy owing to their availability compared to cadaveric islets. We used previously published protocols for the derivation of SC-islets^35,36^ to generate pancreatic progenitors (>60% PDX^+^/NKX6.1^+^) that can then be aggregated and further differentiated towards endocrine lineage and islet-like clusters (Fig. 5a, Supp. Figure 5a-c). Immature SC-islets (end of stage 6) were then either kept in suspension culture, immobilized in 1-2 mm-thick alginate slabs, or embedded inside a device connected to a perfusion bioreactor to evaluate their potential to mature inside the devices. As compared to the compact SC-islet morphology observed in perfused devices, static culture of 10,000 SC-islets immobilised in devices without perfusion led to a rugged morphology likely resulting from substantial cell death (Fig. 5b). Conversely, SC-islets encapsulated inside the device were highly viable, with no necrotic core or peripheral dead cells, and a compact morphology without cell detachment observed via hematoxylin and eosin staining (Fig. 5c-d). Dithizone, a dye staining the insulin granules via zinc specific binding, showed strong insulin presence in the perfused SC-islets versus the static control (Fig. 5e). These data confirm the important advantages of having convective forces increasing oxygen and nutrients transfer for the survival of the SC-islets in microencapsulated devices.

**Fig 5.**
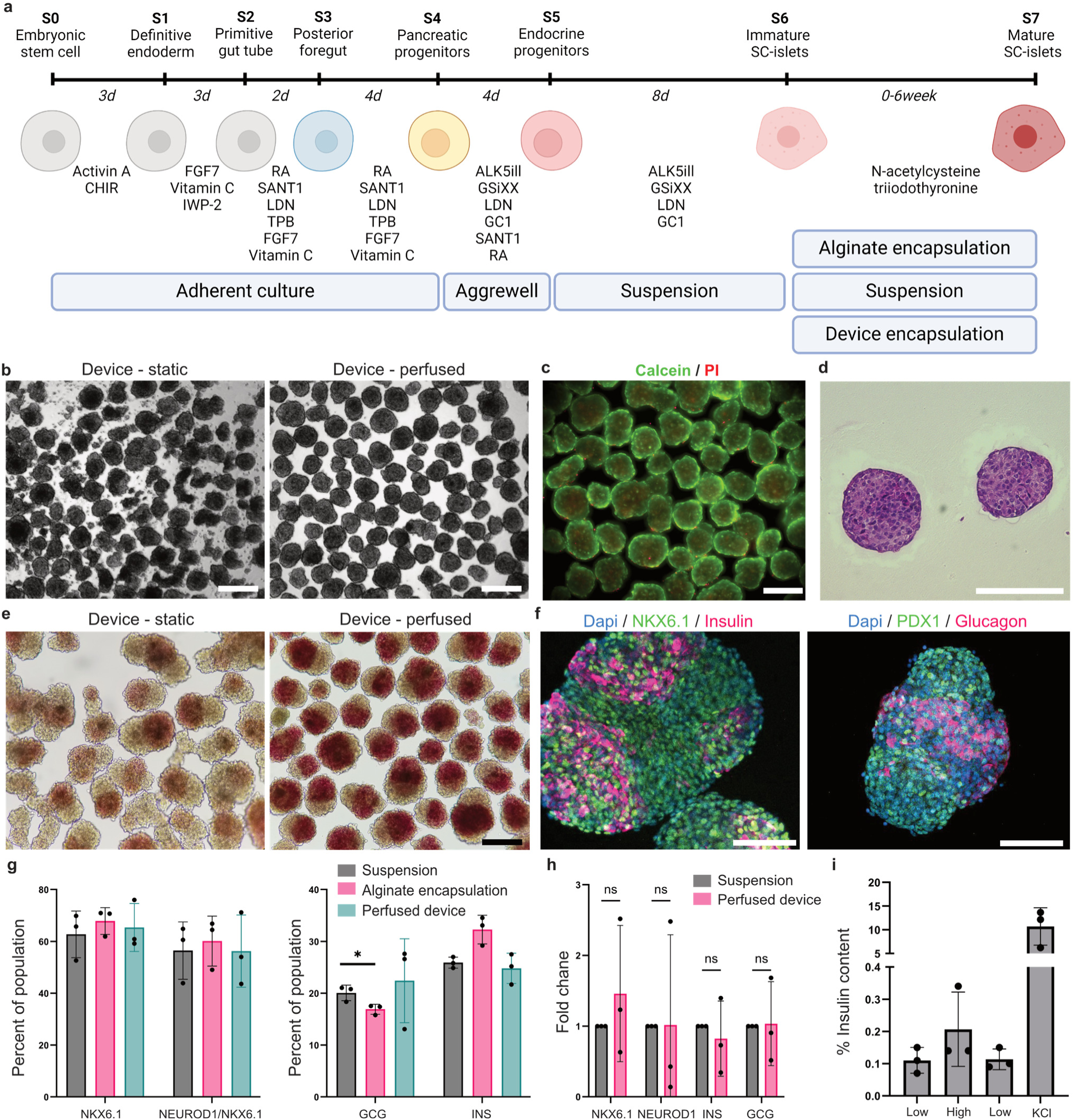
Survival and maturation of stem cell-derived islets during *in vitro* perfusion. **a**, Illustration of the protocol used to differentiate stem cell towards SC-islets. Illustration made with Bio-render. **b**, Representative brightfield images showing SC-islets after 10 days of culture (Stage 7) inside the device either in static or perfused condition. Scale bars, 200 µm. **c**, Representative fluorescent images of live/dead staining of the pseudoislets after perfusion, with Calcein AM (green) and propidium iodide (red). Scale bars, 200 µm. **d**, Hematoxylin and Eosin staining of the SC-islets after 10 days of perfusion. Scale bars, 200 µm. **e**, Representative dithizone staining (crimson red) of the SC-islets after 10 days of culture. Scale bars, 200 µm. **f**, Representative fluorescent images of SC-islets after maturation in the device. Cells are labelled with dapi (blue), NKX6.1 or PDX1 (green) and insulin or glucagon (pink). Scale bars, 100 µm. **g**, Flow cytometry results showing the percentage of the population expressing key pancreatic and endocrine markers. Mean and standard deviation are shown for 3 independent devices with three independent differentiations. *p < 0.05, determined by two-way ANOVA with Tukey’s multiple comparisons test. **h**, Gene expression of the SC-islets after 10 days of perfusion. Mean and standard deviation are shown for 3 independent devices with three independent differentiations. n=3 technical replicates per point. ns=non-significant, determined by a two-way ANOVA with Šídák’s multiple comparisons test. **i**, Insulin secretion in response to changes in glucose concentration or addition of KCl, normalized to the total insulin content. Mean and standard deviation for three independent differentiations with 3 independent devices are shown.

Even though beta cell differentiation protocols have been improved in recent years^36–38^, they generally do not reach the same level of control on glucose sensing and insulin than primary human islets. Interestingly, SC-islets implanted *in vivo* can continue to mature to yield beta cells and other endocrine cells that are more similar to their *in vivo* counterparts. Using SC-islets as a source of insulin producing cells for transplantation thus require cells to be able to mature inside the device after transplantation. To further evaluate the potential for immature SC-islets to mature in inside our devices, we analyzed the endocrine differentiation and maturation during 10 days of perfusion with stage 7 medium. Expression of insulin and glucagon after perfusion was observed via immunofluorescence (Fig. 5f). Flow cytometry and gene expression data showed comparable differentiation efficiency for the perfused device compared to suspension and thin alginate slabs (Fig. 5g-h). SC-islets were also able to respond to step change in glucose concentration and potassium chloride treatment by increasing their insulin secretion (Fig. 5i). Overall, these results demonstrate the feasibility of encapsulating SC-islets inside the device and allowing their maturation during *in vitro* perfusion.

### Transplantation in pigs

Finally, acute safety and patency of cell-free devices was evaluated via one-week device implantation. To assess blood mediated inflammation and cellular infiltration, pigs were not immunosuppressed. Anticoagulation prophylaxis was administrated during and after the surgery, as would be done in humans to prevent complications during the acute phase after transplantation. To facilitate implantation directly as an iliac arteriovenous shunt (Fig. 6a), the devices were modified to include a bend at the exit of the cellular compartment (Fig. 6b). Implantation as a shunt was elected as opposed to an interposition graft to allow hindlimb perfusion in case of device occlusion. One animal received a commercial expanded polytetrafluoroethylene graft (ePTFE) and 6 animals received a device filled with alginate (4 cell-free device and 2 devices with SC-islets). Both the control graft and device were transplanted without acute complication and all seven animals survived until the one-week endpoint (Fig. 6c). Contrary to the ePTFE graft, devices were free of external fibrous encapsulation upon retrieval, confirming the suitability of the material for *in vivo* applications (Fig. 6c). Out of the 4 cell-free devices, graft kinking happened in two devices, preventing blood flow and leading to device clogging. The remaining four devices (two with and two without cells) were patent and had no obvious narrowing of the internal channel, as shown by Doppler ultrasound imaging (Fig. 6d). Pulsatile flow could clearly be seen after implantation (Supp. Movie 2) and prior to removal during Doppler imaging (Fig. 6e). Similar to the ePTFE graft, histological analysis of the devices confirmed limited interaction with blood on the luminal side of the polyurethane graft, with no evidence of thrombi formation within the device lumen or the connecting loop channel (Fig. 6f). Masson Trichrome staining also showed limited internal collagen deposition in the channel, with only small amount of coagulated material in between the alginate and the polyurethane graft (Fig. 6g). Despite the very stringent xenograft model, SC-islets could be identified within the sections (Fig. 6h). Spaces between the gel and SC-islets visible in Hematoxylin and Eosin stains may indicate either some level of cell death or alginate shrinking during fixation (Fig. 6i), but endogenous reporter expression of both insulin and glucagon could be seen in surviving SC-islets (Fig. 6j). Overall, these results nevertheless establish proof-of-concept of short-term human SC-islet xenograft survival in a non-immunosuppressed large animal model. These data confirmed graft patency and safety in the acute phase post-transplant, paving the way for further preclinical studies with this new convection-enhanced device.

**Fig 6.**
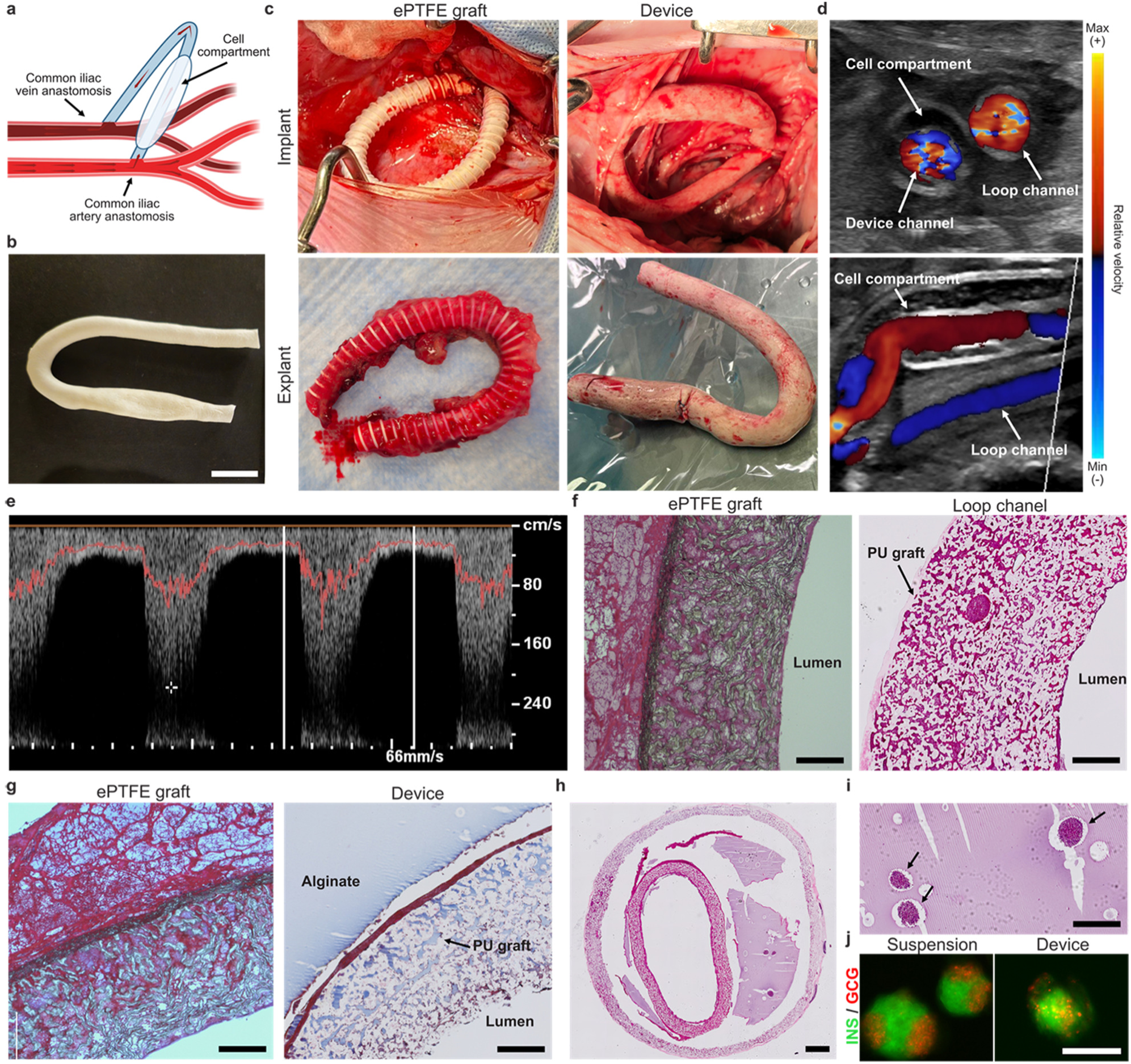
Transplantation of the devices in hybrid pigs. **a**, Illustration of the strategy for device implantation as an arteriovenous shunt between the common iliac artery and vein. **b**, Photographs showing the modified device with a cell compartment on one side and an additional loop for facilitating implantation. Scale bars, 2 cm. **c**, Representative images of the ePTFE prosthesis control (n=1) and device (n=6) at the time of implantation and one week after transplantation as an arteriovenous shunt. **d**, Representative color Doppler ultrasound images of the devices showing the U-shaped loop either perpendicular (top) or parallel (bottom) to the blood flow. **e**, Representative Doppler ultrasound sonographs showing blood velocity through the device. **f**, Hematoxilyn and Eosin staining of the ePTFE control graft and implanted device after 1 week of transplantation. Scale bars, 200 µm. **g**, Masson’s trichrome staining of the ePTFE control graft and implanted device after 1 week of transplantation. Scale bars, 200 µm. **h**, Representative Hematoxilyn and Eosin staining showing cross-section of the device through the polyurethane and cell compartment. Scale bars, 1 mm. **i**, Representative Hematoxilyn and Eosin staining showing SC-islets (black arrow) within the alginate compartment. Scale bars, 200 µm. **j**, Fluorescent images showing glucagon and insulin expression from SC-islets derived from INS^EGFP^GCG^mScarlet^ H1 line. Scale bar, 200 µm.

## DISCUSSION

We present a novel method to fabricate complex vascular grafts within multicompartment devices, and its application towards bioartificial pancreas engineering. This strategy can help overcome critical limitations in diabetes cellular therapy, and artificial endocrine tissue engineering more broadly. It can create complex vascular lattices within a cell-laden hydrogel loading compartment, which can be manufactured, sterilized, shipped and loaded with the intended cell product at the clinical site right before direct anastomosis to the host circulation.

For diabetes cellular therapy, this approach could present significant advantages over existing transplantation approaches. Contrary to other macroencapsulation strategies such as pouches^10,39^, sheets^40^, or oxygenation chambers^15^, the internal vascular lattice can provide immediate blood flow through the graft. This can reduce the significant islet losses that occur during the anoxic period post-transplantation observed even in the hepatic site, thereby potentially reducing cell doses needed to reach therapeutic effects. It has been estimated that the surface area to avoid oxygen limitations to islet survival and function in macroencapsulation relying solely on diffusion is on the order of a meter square^9^. Our proposed design addresses this challenge through 3D printing to create vascular lattices that provide oxygen and insulin transport via bulk motion to the external compartment where transport occurs via diffusion. Because of the higher surface area created by the vascular lattice, as well as the arterial oxygen tension that can be provided at the interface, our devices can accommodate higher number of cells than diffusion-based devices of the same islet compartment volume. According to numerical models and experimental findings, our current 9-channel devices could accommodate 70 × 10^6^ SC-islets while maintaining adequate oxygenation in most of the 7.2 mL cell and hydrogel loading chamber (Supp. Figure 6). This corresponds to ∼40,000 islet equivalents (IEQ), as compared to ∼8,000 IEQ/kg required to consistently achieve insulin independence with hepatic islet transplants^41^. However, it has been estimated that over half of the islets fail to engraft in the liver, with 4,000 IEQ/kg (500 × 10^6^ cells) likely sufficient if most SC-islets survive^42,43^. To meet these requirements, devices with higher channel numbers (e.g. 3-fold more channel surface area) and increased length (e.g. 20 cm instead of 5 cm current) could be developed by combining numerical optimization with the unique capacity to generate complex pre-determined vascular networks shown here. As compared to pouch-based devices such as ViaCyte’s PEC-Encap® which was re-engineered to allow graft vascularization (PEC-Direct®), the proposed design maintains a barrier between the graft and the recipient while still permitting internal vascularization^10^. This can provide the “best of both worlds”, avoiding direct contact between immune cells and the islets, while maintaining adequate oxygen, nutrient, waste product and insulin diffusion rates for graft function. Microencapsulation studies in rodents and primates have shown that long-term islet survival and glycemic control does not require direct islet vascularization^33,44^. Our macroencapsulation device presents advantages like microencapsulation in terms of higher surface area/volume ratios than single compartment devices, but also localizes the graft in a well-defined site. Even with emerging gene editing strategies to create grafts that evade immune detection or rejection^45,46^, devices that hinder graft cell escape improve the safety profile of stem cell-based grafts.

Convection enhanced devices have been proposed, and proof-of-concept studies and computational models suggest that they increase the cell-loading capacity and compactness of the device while decreasing cellular necrosis^17^. Cell-impermeable convection based macroencapsulation devices developed to date either require an external pump^17^ or connection to a commercial graft that will then be sutured to blood vessels^19,21,47^. The latter approach is however susceptible to rupture due to material interfaces that can be prone to fibrosis, thrombosis or mechanical fatigue. Humacyte recently described an inter-vascular device with a single-channel acellular tissue engineered vessel, which would be expected to encounter the same initial oxygenation limitations as our single-channel prototypes^21^. To overcome these challenges, we hypothesized that an ideal intervascular device would allow for the fabrication of complex geometry of the internal vascular network while being made from only one material that can be directly anastomose to the host blood vessels. Although interesting approaches to hierarchical vascular grafts have been developed^25,48^, they would require complex casings or rely on fusion processes to incorporate them into macroencapsulation devices. Alternatively, various 3D printing approaches have been applied to create hierarchical vascular networks within centimeter-scale cell-laden hydrogels^49,50^. However, these cell-laden hydrogels are not readily anastomosed to blood vessels without multiple material interfaces that can diminish the long-term patency of the device^2,5,24^.

This work demonstrates the feasibility of engineering convection enhanced devices that can be connected to flow loops *in vitro* for long-term bioartificial organ culture or anastomosed to vasculature in large animal models. Our experimental findings with beta cell lines as well as stem cell-derived islets demonstrate the advantages of the vascular lattice device over bulk cell immobilization strategies, as predicted by mass transport models. Perfusion in the devices leads to rapid insulin release into the circulation, combined with large cell viability rings seen around the parallelized vascular network – features that will be critical for translation of the technology to patients. Using a custom-made bioreactor, we also demonstrate that SC-islets can survive and mature efficiently inside the device. This is important since maturation of SC-islets is incomplete *in vitro,* and it can take several weeks to months *in vivo* before transplanted cells reach gene expression levels comparable to human islets^51^. Even in a challenging xenogeneic setting *in vivo*, some SC-islets survived after one week of transplantation. These results are encouraging since allogeneic cell sources such as SC-islets and human islets intended for clinical use would likely encounter a less stringent immune rejection context. We demonstrated that the devices can also be implanted in heparin-treated pigs as an arteriovenous shunt without acute complications and remain patent for at least a week with limited fibrosis noted on the external pouch or in the channel lumen. Despite these strong proof-of-concepts from our first device prototypes, there are some limitations that warrant further optimisation. Currently, the fabrication process results in vascular prostheses that are susceptible to kinking when challenged by surgical intervention followed by internal organ movements in the animal. To address this issues, future device iterations can be reinforced using modified 3D-printed lattices, combination with other technologies such as electrospinning or melt electrowriting, or by providing external support to the device when it is necessary to incorporate curvature for implantation. Although the devices performed well during a short implantation window, longitudinal studies will have to be conducted to confirmed patency and absence of thrombogenic events for multiple months. We also expect that optimization of the 3D printed vascular network geometry will be critical to optimize hemodynamic flow within the channels.

We can leverage the modularity and flexibility of our approach to independently improve device geometry, immunoprotection, cell sourcing and vascular interface. The fabrication and delivery process are compatible with various cell sources (human islets, SC-islets, porcine islets) and hydrogels that could further prevent physical immune interactions or modulate the immune system to create a tolerogenic environment for the graft^52^. Emerging approaches like the engineering of immunoevasive SC-islets^45,53^ could also be combined with our device when paired with a hydrogel promoting vascularization within the device. This approach would lead to lower overall device sizing requirements due to secondary vascularization via microvasculature, potentially even allowing long-term device function even if long-term internal flow occlusion occurs.

To conclude, we present a new fabrication technique that allows for the design of an intervascular macroencapsulation device with controllable geometry of the internal vascular network. This study represents an important step forward in the development of intervascular devices for the treatment of type one diabetes, where we merge concepts of complex vascular network fabrication and intervascular macroencapsulation devices. This technology represents a promising direction for long-term implantation in diabetic pig and could be compatible with cell therapy delivery in human.

## METHODS

### Vascular graft and encapsulation device fabrication

#### Sacrificial carbohydrate molding (single channel and pouch)

Plastic molds were first designed using AutoCAD (Autodesk) and printed using a TAZ workhorse 3D printer (Lulzbot) with Cura LE v3.6 software and black polyLite PLA (Polymaker). Polydimethylsiloxane (PDMS, Dow Chemical Company) was prepared by mixing the elastomer with the curing agent (10% v/v) and was cast over the plastic molds to generate inverted molds for the sugar templates. PDMS was cured overnight in an oven at 50°C. The sugar glass mixture was prepared by mixing 113 g of sucrose (Fisher) and 75 mL of distilled water in a 500 mL beaker, followed by heating on a digital hotplate stirrer with a built-in feedback temperature control system (Scilogex, MS7-H550-Pro) until the temperature of the liquid reached 175°C. The solution was stirred at 400 rpm with a 1-inch magnetic stir bar during the heating process. The liquid sugar solution was then poured into the preheated (100°C) PDMS molds and solidified at room temperature for 20-30 min before demolding. Sugar constructs with small imperfections were polished using a sharp scalpel blade.

#### 3D printing of sugar templates (9-channel graft)

Immediately after preparation, the molten carbohydrate ink (78 g sucrose/50 mL water) was loaded into 20-mL glass syringes (Fortuna Brand), which were then inserted into a modified Airwolf AW3D XL printer. This printer was equipped with a previously developed printer head^24^, and two cooling air jets (7.5 PSI) were placed on opposite sides of the nozzle outlet. To produce optimal printability^54^, minimal additional caramelization, and rapid solidification of the original material, the following parameters were used: printer head with 2.0 mm extrusion nozzle, volumetric flow rate of 0.123 g/min, translational speed of 15 mm/min, printing head and nozzle temperature of 92°C and 85°C, respectively. Custom G-codes were written to produce the sacrificial sugar network. Each channel with 9 branches took 80 min to print, and three sugar templates could be printed with one preparation of sugar. The sacrificial templates were then stored at 4°C in airtight packaging with moisture-absorbing silica gel, shipped from Québec to Montreal (3h driving distance) and were <u>used</u> within three months.

#### Dip coating and fabrication process^55^

3D printed or molded sugar templates were dip coated in a polycarbonate polyurethane solution composed of 40 mL of ChronoFlex® AR 22% (AdvanSource Biomaterials Corp), 25 mL of N,N-dimethylformamide (DMF, Sigma-Aldrich) and 12,93 g of sodium chloride crystals (NaCl, Sigma). For optimization of the coating using factorial design as described below, the salt concentration was varied to obtain 1:2, 2:2 and 3:2 ratio NaCl/dry polymer w/w (corresponding to 33%, 50% and 60% of porogen), without modifying the ratio of DMF to ChronoFlex. Sugar lattices were coated 3 to 5 times with 2h drying intervals in a fume hood and dried overnight before being transferred to reverse osmosis water to dissolve the sugar and salt content. For device fabrication, the coated sugar networks were placed in a two-piece PDMS mold providing the geometry of the external pouch, and filled with a sugar preparation cooled down to around 80°C. The external pouch coating was made using the same methodology as described above, with three dip coatings in the polymer and salt mixture and one dip coating in the mixture without porogen. The last coating was made without including salt in the solution to create a smooth finish to create a barrier to avoid fluid outflow from the device.

### Graft and device characterization

#### Full factorial design

Polyurethane grafts were fabricated as explained above with 6 mm diameter sugar rods. Design of experiments was applied using JMP® Statistical Software (SAS Institute, version 17). A 3 × 3 full factorial design was applied to examine the effect of salt crystal size (S, 3 levels: S=45 µm, S= 75µm, S=150 µm) and concentration (C, 3 levels: 33%, 50%, 60%, corresponding to salt/polymer weight ratios of 1:2, 2:2 and 3:2) on two output parameters - porosity and longitudinal tensile strength of the grafts. Measures for each of these 9 conditions were independently replicated 5 times. A linear model was applied to estimate the main and first order interactions of S and C. Using the surface profiler function, surface response curves were generated for both the porosity and the tensile strength, and their predicted formula were extracted. We then used the custom profiler to generate a desirability function that would maximize both the tensile strength and the porosity (30% vs 70% weight respectively). Maximization of this desirability function was then used as the criteria for choosing the salt size and concentration.

#### Physical property measurements

To measure the diameter expansion, brightfield microscope (VWR) pictures of circumferential sections of the graft were taken before and after hydration. Using Fiji/ImageJ (NIH), graft thickness and internal diameter were measured for four cross-sections for each graft and external diameter expansion was calculated as a percentage of the difference of the diameter in wet and dry conditions over the diameter of the dry cross-section. Polymer deposition was measured by dividing the weight of the dry prosthesis by their length.

#### Porosity characterization

After soaking in distilled water for 36 to 48 h, the grafts were gently shaken to remove excess water and dried overnight at 55°C before being weighted (W_dry_). To measure the volumetric porosity in dry conditions, the grafts were then submerged in hexane for 1h, shaken to remove excess liquid and weighed again (W_hex_). The volumetric porosity was then calculated using the following equation, with known hexane density and experimentally measured ChronoFlex density:

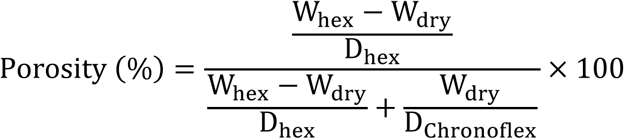

#### Permeability

Grafts were placed in our custom-made vascular bioreactor (Supp. Figure 4) with the external compartment filled with water and a flow loop attached to a peristaltic pump (Masterflex). Next, 20 kDa FITC-dextran (0.4 mg/mL in water) and circulated through the graft at 40 mL/min. The set up was arranged so that the bioreactor and graft could be imaged every 12 s in a small tissue imager (IVIS Lumina LT, 1s exposure, medium binning). The images were first exported in logarithmic scale from the IVIS software and analyzed using Fiji/ImageJ (NIH) to produce stills and videos of the dextran diffusion.

#### Longitudinal tensile strength (LTS)

For the graft optimization, measurements were performed on a CellScale (Univert) according to ISO 7198 for cardiovascular implants and extracorporeal systems. Grafts were cut perpendicular to the long axis to obtain 2 cm segments and strained parallel to the long axis at 60 mm/min until failure. The highest force was recorded as longitudinal tensile strength. LTS data comparing the effect of layer coating and vascular grafts versus single-channel graft with external compartment (devices) were collected using a universal testing machine (Instron 5965, Instron Inc.) using the same parameters and protocol. Distance between grips were set at 50 mm and devices were made thicker at the extremities with multiple coating only at the extremities (∼2 cm) to prevent slipping during testing.

#### Suture retention strength (SRS)

SRS data was collected using a universal testing machine (Instron 5965) according to ISO 7198. Vascular prostheses were cut perpendicular to the longitudinal axis in 2 cm long segments. A 7.0 prolene suture (Ethicon) was pierced through the prosthesis wall at 2 mm from the edge with the two sides of the suture looped together with duct tape to prevent slippage from the Instron grip. The suture loop and the untouched end of the graft were each attached to a separate clamp in the universal testing machine. The sutures were pulled at 60 mm/min until rupture of the graft wall and peak force was recorded as the suture retention strength.

#### Pressurized internal radius and burst pressure

Grafts were cut perpendicular to the longitudinal axis to make 5 cm segments and attached to our custom bioreactor connectors using tie wraps. A closure was affixed to the outlet to allow pressure build-up inside the graft. A latex sleeve (Long party balloon) was affixed to surround the graft and prevent the flow of water through the pores. The sleeve was pre-stretched manually and assumed to have negligeable resistance. The open end of the bioreactor was connected to a syringe pump (Harvard Apparatus) and a pressure monitor (World Precision Instrument). Water was first infused at a rate of 0.5 mL/min to measure the internal radius under known pressure until 6 psi, then effused until the pressure fell to zero and then infused again at 25 mL/min until burst (average of 1.5 psi/second). A Sony Cyber-shot WX-100 was used to acquire images at each desired pressure (50, 100, 200, 300 mmHg) and videos of the burst experiments. Image processing and measurements were performed manually using Fiji/ImageJ (NIH) to measure the radial expansion for each graft whereas thickness was measured on 1-2 mm length graft cross-sections imaged under phase contrast microscopy (VWR). Burst pressure was defined as the highest pressure recorded on the pressure monitor during the burst experiment. Assuming an incompressible wall, compliance was calculated as follows:

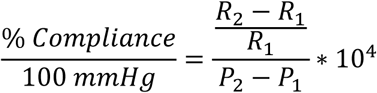

Where P = pressure, R_1_ = radius at P_1_ and R_2_ = radius at P_2_.

#### Biaxial tensile tests

Grafts were sterilized by either immersion in 70% ethanol for 2 h or by autoclaving in a humid vapor cycle for 30 min. Grafts were then cut open and trimmed into 1.5 cm × 1.5 cm squares which were attached to an ElectroForce TestBench (TA Instruments, New Castle, Delaware, USA) with 4-0 hooked silk sutures in a bath of reverse osmosis water at 37 °C. Each sample underwent cyclic loading–unloading to 40% strain at 0.4 mm/s for 8 preconditioning cycles followed by four data cycles at a displacement rate of 0.1 mm/s. Strain measurements for the last three were recorded and averaged to calculate the apparent modulus using the tangent at 50% of strain.

#### Platelet isolation and adhesion

Platelets were isolated from human blood following a previously described protocol^56^. Briefly, whole blood was collected in sodium citrate tubes following a protocol approved by the McGill Institutional Review Board (Study Number A06-M33-15A). The whole blood was centrifuged at 350 × g for 20 min to collect the platelet rich plasma (50% top volume fraction, PRP). PRP was diluted in calcium-free Hanks’ Balanced Salt Solution (HBSS, ThermoFisher) supplemented with 4 mM EDTA (ThermoFisher) buffered to pH 6.4 and centrifuged again at 350 × g for 20 min to recover the platelets. Platelets were then resuspended in HBSS-EDTA and distributed at a density of 2.5 ×10^7^ platelets/cm^2^ over 0.5 cm^2^ sections of prosthesis held down by a PDMS ring to prevent the graft from floating. A positive control for platelet activation was obtained through platelets exposed to 100 µM ADP (Sigma) for 1 min at 37°C. Samples were incubated for 2 h at 37°C before being gently washed in phosphate buffered saline without calcium and magnesium (PBS) and processed using a LDH cytotoxicity assay kit (Cayman) following the recommended protocol. Absorbance was measured using a Benchmark Plus microplate reader (Bio-Rad).

#### Endothelial cell adhesion

Human umbilical vein endothelial cells (HUVECs) were seeded at 1 ×10^4^ cells/cm^2^ over 0.5 cm^2^ sections of prosthesis held down by a PDMS ring inside 24-well plates with 1.5 mL of endothelial cell growth medium-2 (Lonza). After 2 h of adhesion, half of the grafts were gently washed and transferred to fresh medium containing WST-8 and incubated for 1h. The other half of the grafts were kept in culture for 4 days, gently washed and transferred to fresh medium containing WST-8 and incubated for 1h. For each condition, 100 µL of supernatant was transferred to a 96-well plate and absorbance (450 nm) was measured using a Benchmark Plus microplate reader (Bio-Rad).

### Microscopy and image acquisition

#### Scanning electron microscopy (SEM)

Graft structure was observed using a variable pressure scanning electron microscope (VP-SEM, Hitachi SU-3500). Grafts were dehydrated and cut into 0.5 cm × 0.5 cm segments before imaging the internal surface. Brief immersion in liquid nitrogen was followed by fracture of the prosthesis to image the cross-section topography and porosity. SEM pressure was set at 40 Pa with accelerating voltage between 4 and 10 kV depending on the magnification. To image adhesion of platelets, grafts were gently washed with PBS and fixed using 2.5% glutaraldehyde in PBS for 2h at room temperature. The samples were then dehydrated in successive ethanol-water mixtures (50%, 70%, 80%, 90%, 95%, 95%, 100%, 100 vol%) for 15 min each, and dried overnight at room temperature before imaging as described above.

#### Immunostaining

Grafts and tissues were rinsed twice with PBS and fixed with 4% paraformaldehyde (Thermo Fisher Scientific) in PBS for 15-20 min at room temperature. Following fixation, tissues were permeabilized with 0.2% Triton X-100 in PBS for 1 h at room temperature, followed by blocking with protein block solution (Dako) for 3 h at room temperature. The samples were then incubated overnight in humidified chamber at 4 °C in the presence of antibodies against VE-cadherin (1:200, R&D, catalog no. MAB9381), NKX6.1 (1:100, DSHB, catalog no. F55A10), PDX1 (1:200, R&D, catalog no.MAB2419), insulin (1:200, Invitrogen, catalog no. 701265) or glucagon (1:200, Invitrogen, catalog no. 14-9743-82) in antibody diluent (Dako). After washing three times in PBS for 1h at room temperature, samples were incubated overnight at 4°C with Alexa 647 donkey-α-rabbit (1:500 in blocking solution; Invitrogen) or Alexa 647 donkey-α-mouse (1:500 in blocking solution; Invitrogen) secondary antibodies in antibody diluent containing DAPI (1:2,000). After at least 3 x 1h washes in PBS, samples were mounted in fluoromount-G (Thermo Fisher Scientific) before imaging. Fixed samples were imaged with an inverted confocal microscope (INVERT Zeiss Axio Observer Z1) equipped with ×10/0.30 NA and ×20/0.80 NA air objectives; 405-, 488- and 555-nm lasers and controlled by ZEN 2009 imaging software (Zeiss). Image processing was performed using Fiji/ImageJ (NIH), with brightness and contrast adjustments only.

#### Live/Dead staining

Thin vertical slices of the inlet or center of the device were cut using a razor blade and stained for 20 min at 37 °C using a live/dead solution with final concentrations consisting of 18.9 µg/mL propidium iodide (Fisher Scientific) and 1.1 µg/mL Calcein AM (Fisher Scientific) diluted in culture medium. Single channel devices and SC-islet images were acquired either using an IX81 Olympus Microscope using a FITC filter cube (Ex: 482/35 | Em: 536/40) and a Texas Red filter cube (Ex: 525/40 | Em: 585/40). Large-scale device images were acquired on a Zeiss Axio Observer fully automated inverted microscope with ZEN 2009 imaging software (Zeiss). Image processing was performed using Fiji/ImageJ (NIH), with brightness and contrast adjustments only.

#### Histology

Samples were fixed overnight in a modified Bouin’s solution (150 mL of water, 50 mL formaldehyde, 10 mL of glacial acetic acid), transferred to 70% ethanol and sent to the Institut de recherche Clinique de Montréal (IRCM) for paraffin embedding, sectioning, deparaffinization and staining. Sections of 6-μm thickness were taken orthogonal to the perfusion channel and stained with either Hematoxylin and Eosin or Masson’s Trichrome solution. Imaging was performed using a Zeiss AxioImager Z2 automated upright microscope and Zen Pro acquisition and analysis software.

### Cell culture and characterization

#### Cell lines culture

Mouse insulinoma 6 (MIN6) cells (kindly donated by Jun-ichi Miyazaki, Osaka University, Japan)^57^ were maintained in DMEM high glucose supplemented with 10% FBS, L-glutamine (1x), Penicillin-streptomycin (1x) (all from Invitrogen) and β-mercaptoethanol (Sigma), and used between passage 25 and 40. Unlabelled HUVECs or HUVEC with a constitutive green fluorescent protein (GFP) reporter (CMV-GFP HUVEC, ATCC) were cultured in EGM-2 Bulletkit medium (Lonza) and used before 10 passages. Medium was changed every other day for all cultures. Cells were maintained in cell culture treated polystyrene plates (Sarstedt – red) at 0.133 mL/cm^2^ medium volume.

#### Pancreatic differentiation

For the *in vitro* studies, human embryonic stem cells (WA01 H1) were obtained from WiCell and used between passages 30 and 40. For the *in vivo* study in pigs, H1 INS^EGFP^GCG^mScarlet^ line was provided by Dr. Francis Lynn^58^. Culture surfaces consisted in Sarsted (red) plates coated for 120 min at room temperature with hESC-qualified Matrigel for routine maintenance, or growth factor reduced Matrigel for differentiation. The hESCs were cultured in mTeSR1 medium (STEMCELL Technologies) and passaged by washing in Ca^2+^ and Mg^2+^ free PBS, incubating for 4 min in 50 µM EDTA in PBS, replacing the EDTA solution by medium, removing clumps from surfaces, triturating gently with a 10-mL pipet to obtain clumps, and re-seeding at 1:3 to 1:30 dilution. At the start of the differentiation, cells were dissociated using TrypLE (Invitrogen) and seeded as single cells (density of 1.35 × 10^5^ cells/cm^2^) in mTeSR1 medium supplemented with 10 µM Rho-Associated kinase inhibitor (Stem Cell Technologies). The pancreatic differentiation was done following previously described protocols^36,38,59^, with some modifications. Medium was changed every day for stage 1 to 6 and every other day during stage 7. Complete media formulations and growth factors are supplied in Supplementary Table 2. After the third day of Stage 4 culture, cells were detached using TrypLE and seeded in AggreWell400 plates (STEMCELL Technologies) to obtain 1000 cells/aggregates. During stage 5, cells were fed directly in the AggreWells, with 70% medium change daily. Aggregates were transferred to ultralow attachment plates (Corning) at the beginning of stage 6 and placed on a Celltron orbital shaker (Infors HT) at 100 rpm. At the beginning of stage 7, aggregates were either kept in suspension, encapsulated in alginate or encapsulated inside the devices and cultured for 10 more days with medium change every other day.

#### Flow cytometry

Monolayer and SC-islets were dissociated using TrypLE for 5–10 min at 37 °C and resuspended in PBS supplemented with 2% FBS. Cells were incubated for 20 minutes with fixable viability dye (L34963, Life Technologies) and washed one before being fixed and permeabilized using Cytofix/Cytoperm (BD Biosciences) for 10 min. Cells were washed in PBS and resuspended in Perm/Wash buffer (BD Biosciences). Cells were then incubated with conjugated antibodies for 30 minutes: SOX17-PE (561591, BD Biosciences), PDX1-PE (562161, BD Biosciences), NKX6.1-AF647 (563338, BD Biosciences), NEUROD1-PE (563001, BD Biosciences), glucagon-PE (565860, BD Biosciences), c-peptide-AF647 (565831, BD Biosciences), AlexaFluor® 647 mouse IgG1 K Isotype Control (557732, BD Biosciences), PE mouse IgG1 K Isotype Control (554680, BD Biosciences). They were then washed with Perm/Wash buffer and resuspend in PBS with 2% FBS for analysis. The cells were run on a BD FACSCanto II cytometer (BD Biosciences) and analyzed with FlowJo v.10 (BD Biosciences). For the gating strategy, debris were removed from analysis using FSC/SSC gating, and dead cells were removed by gating the fluorescent cells having incorporated the fixable viability dye. Isotype control was used to gate the antibody-stained cells.

#### Quantitative RT-PCR

To extract total RNA, RNeasy® Mini Kit (Qiagen) was used according to the manufacturer’s protocol. The iScript™ gDNA Clear cDNA Synthesis kit (Bio-Rad) was used to generate cDNA. The qPCR reactions were performed using SsoFast™ EvaGreen® Supermix (Bio-Rad) on a ViiA7 (ThermoFisher) according to the manufacturer’s protocol. Data were normalized to the housekeeping gene NFX1 and data is presented in relation to either the suspension protocol (perfusion experiments) or human islets (protocol characterization). Primers were synthesized by Integrated DNA Technologies Inc. and are provided in the Supplementary Table 3.

### Device perfusion and bioreactor operation

#### Device preparation and perfusion

Devices were extensively cleaned with distilled water (3 times 24 hours in 4L beaker) and installed into custom-made vascular bioreactor (Supplementary Figure 4) using autoclaved resistant tie-wraps. The vascular bioreactor was then assembled and connected to the perfusion tubing assembly. The bioreactor’s ports were loosened and the whole set-up was autoclaved in a humid vapor cycle for 30 min. For MIN6 encapsulation, internal gelation of Protanal 5% was used. Briefly, alginate 6% was prepared by dissolving the alginate powder in HEPES solution (10 mM 4-(2-hydroxyethyl)-1-piperazineethanesulfonic acid (HEPES), 170 mM NaCl – all from Sigma, pH 7.4) and mixed with CaCO_3_ (VWR) and Glucono-δ-lactone (Sigma). Proportions were adjusted to yield a hydrogel solution with final concentrations of 5% alginate, 30 mM CaCO_3_ and 0.6 mM Glucono-δ-lactone with 20 or 40 million MIN6 cells per mL. After cell addition to the mix, the solution was slowly injected inside the device, and perfusion was started after 15-20 min of gelation. For SC-islets encapsulation, cell aggregates were mixed with ultrapure alginate (50:50 mixture of ultrapure LVM alginate and MVG alginate (Novamatrix) to produce 5% alginate hydrogel solution. The solution was injected inside the device using 18G sharp end needle through the external compartment near the inlet of the device. To achieve external gelation via diffusion of divalent cations from the graft into the external alginate-filled compartment, a concentrated calcium chloride (Sigma) solution (10 mM HEPES, 75 mM NaCl, 100 mM CaCl_2_, pH 7.4) was poured in the external bioreactor compartment slowly perfused via the graft channel inlet for 15 min to allow for hydrogel gelation. The CaCl_2_ solution was removed from circulation, replaced with stage 7 medium supplemented with 2 mM CaCl_2_ and set-up inside the incubator under perfusion conditions. Perfusion of culture medium was established using a peristaltic pump (Ismatech) set-up inside an incubator at 37°C and CO_2_ was maintained at 5%. Pump tubing (VWR Internation, MFLX95723-48) with 2.79 mm internal diameter was connected to silicone tubing (VWR Internation, MFLX07625-46) to facilitate gas equilibration. A culture medium flow rate of 25 mL/min (1-channel device) or 40 mL/min (9-channel device) was applied except for oxygen consumption and insulin secretion experiments. The medium was drawn from a 50-mL reservoir of culture, and medium was manually exchanged every two days by bringing the set-up under a biosafety cabinet.

#### Glucose-stimulated insulin secretion

Devices were perfused in an open-loop at 1.5 mL/min for 2 h with Krebs buffer supplemented with 2.8 mM glucose to equilibrate to basal glucose in the system. Krebs buffer contained 129 mM sodium chloride (Sigma), 4.7 mM potassium chloride (Sigma), 1.2 mM MgSO_4_ (Sigma), 10 mM HEPES (Fisher), 2.5 mM CaCl_2_ dihydrate (Sigma), 5.0 mM NaHCO_3_ (Fisher) and 0.5% w/v bovine serum albumin (Sigma, A3294) in reverse osmosis water. The pH was brough to 7.4 using sodium hydroxide (Fisher). The insulin secretion test was performed by applying sequential Krebs solutions supplemented with 2.8 mM glucose for 30 min, 16.7 mM glucose for 45 min, 2.8 mM glucose for 45 min and 30 mM KCl for 30 min. For the KCl, NaCl concentration was reduced to 104 mM to maintain adequate osmolarity. Samples were taken from every 3-min fraction, quickly centrifuged to remove potential cell impurities and stored at -20°C. For analysis, the thawed samples were processed using an ELISA kit for mouse (Cedarlane, 80-INSMS-E01, with standard curve) or human (Cedarlane, 80-INSHU-E01.1, with standard curve) according to the manufacturer’s instructions. Absorbance (450 nm) was measured using a Benchmark Plus Microplate Spectrophotometer (Bio-Rad).

#### Oxygen measurement

Using the Pyroscience O2Logger software, the oxygen probe signal was adjusted using 2-point calibration at the following conditions: (i) 0% O_2_ using water with an oxygen scavenger (OXCAL, Pyroscience) and (ii) air-saturated water at a set temperature in the software corresponding to the temperature measured with an alcohol thermometer. Prior to acquisition, the bioreactor was first perfused overnight, and oxygen concentration monitoring was done with the probe and bioreactor remaining in the incubator at 37°C and 5% CO_2_. The oxygen probe’s needle was inserted inside the tubing at the exit of the bioreactor and held in place close to the middle of the channel. Oxygen concentration was acquired every minute for the duration of the experiment and each flow rate was maintained for 1 or 2 h before being manually changed to the next setpoint.

#### Finite element modeling of oxygen profiles

SolidWorks models of the single and nine-channel devices were imported into COMSOL Multiphysics 5.5 software. The flow of cell culture medium (μ = 0.69 mPa·s, ρ = 0.993 g/cm³) was modeled with a no-slip condition applied to all internal walls of the devices. A time-dependent laminar flow solver with a relative tolerance of 0.01 was implemented. The fluid (culture medium) was assumed to be incompressible and Newtonian, with constant viscosity. A plug flow was specified at the inlet, with flow rates ranging from 0.75 to 25 mL/min for the single-channel device filled with MIN6 cells, and up to 800 mL/min for the nine-channel device filled with SC-islets. The oxygen concentration within the devices was modeled using the Transport of Diluted Species module. The inflow oxygen concentration was set to 0.198 mol/m³. The single channel perfusable devices with either 40 × 10^6^ cells/mL, 20 × 10^6^ cells/mL and 10 × 10^6^ cells/mL MIN6 and 5 × 10^8^ cells MIN6 cells were simulated as described above for 2 days.

The diffusion coefficient of the polyurethane interior wall was considered, while the outer wall of the device was treated as an impermeable barrier to solve the diffusion model described in Eq. 1.

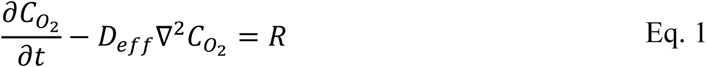

D_eff_ denotes the effective oxygen diffusivity within the beta cell-laden alginate, C_o2_ indicates the radial oxygen concentration, and R signifies the rate of oxygen consumption. As outlined in Eq. 2, we calculated the oxygen diffusivity as a weighted combination of the diffusivity in the alginate matrix (D_alg_) and within the cells (D_cells_), where f represents the volumetric fraction occupied by cells.

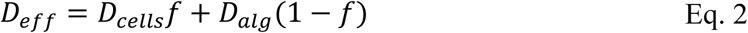

To define R, we use the Monod oxygen consumption kinetics described in Eq. 3.

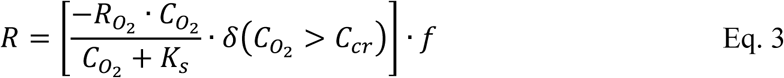

Here, R_o2_stands for the maximum oxygen consumption rate, K_s_ is the Monod coefficient related to oxygen uptake, o represents the Heaviside function and C_cr_ signifies the critical oxygen concentration threshold below which cell death (lack of oxygen consumption) occurs. Table 4 provides an overview of all model parameters.

### Animal experiments

#### Animals

Murine experiments were performed at University of British Columbia following protocol approved by the UBC Animal Care Committee (Protocol A21-0045). NSG mice (NOD.Cg-Prkdcscid Il2rgtm1Wjl/SzJStrain Code: 5557) were obtained from Jackson Laboratories, Bar Harbor, ME. Pig experiments were conducted at the Centre Hospitalier de l’Université de Montréal following protocols approved by Institutional Animal Care Committees at the CHUM and at McGill (Protocol A23009MBp). Pigs from Ferme Triporc Inc. were used. The animals were non-diabetic, immunocompetent pigs weighing 35 to 50 kg.

#### Subcutaneous implantation in mice

The vascular grafts were cut into small rings of 2 mm in length, sterilized in 70% ethanol overnight and thoroughly washed with sterile PBS before implantation. Mice were anaesthetized with inhalable isoflurane and two rings were implanted per animal, one on each side of the back. A small incision was made, and the ring was placed in a subcutaneous pocket made with blunt scissors. The incision was closed with absorbable subcuticular suture (type 5.0). The mice were euthanized to retrieve the devices after 4 weeks. The rings were washed with HEPES buffer, put in modified Bouin’s fixative, and incubated in this fixative overnight. Samples from each device were paraffin embedded and sectioned radially for further staining.

#### Arteriovenous shunt surgical procedures in pigs

The geometry of the device was modified to create a ‘U-shaped’ polyurethane extension to facilitate implanting them as an arteriovenous shunt connecting the right iliac artery and the right iliac vein. Cell-free devices were prepared by loading the devices with alginate the day prior to surgery and were incubated in HEPES buffer + 40 mM CaCl at 4°C. To prepare the cell-laden devices, SC-islets were mixed with alginate and loaded in the devices 2 h before the surgery and stored on ice in Stage 7 medium supplemented with 40 mM CaCl_2_ until implantation. The iliac artery and vein were first mobilized. The iliac vein and artery were clamped followed by the polyurethane extension-to-vein anastomosis and device end-to-artery anastomosis at 45° angle using 6-0 polypropylene sutures ensuring that the device receives blood from the artery and outflow to the vein. The vascular clamps were removed, and the perfusion of the device was confirmed by assessing pulsation in the device. The incision was closed with #1 polydioxanone suture and the skin was closed with 3-0 vicryl sutures. As pre/post-operative care, the pigs were administered antiplatelet drugs (clopidogrel, 75 mg), anticoagulant medication (Apixaban, 0.5 mg/kg), anti-inflammatory drug (meloxicam, 0.3 mg/kg), and gastric ulceration prevention medication (Esomeprazole, 40 mg). The pigs were monitored for appetite, surgical wound healing, and general exhibition of stress or lethargy until study endpoints. Seven to eight days post-transplantation, the pigs were euthanized to retrieve the devices. The devices were excised from the vasculature and perfused with HEPES buffer to remove any residual blood in the channel. The devices were then perfused with modified Bouin’s fixative and incubated in this fixative overnight. Samples from each device were paraffin embedded and sectioned radially for further staining.

#### Doppler imaging

All ultrasound images were acquired on a EPIQ G7 system (Philips, Amsterdam, The Netherlands) using a 12-3 MHZ linear probe. First, B-mode acquisitions were performed to evaluate the localization of the bypass. Then, the patency was assessed using color doppler ultrasound. Duplex ultrasound was performed with an angle of less than 60 degree on the arterial and venous anastomoses, the proximal, mid and distal portion of the bypass. The flow was measured using the Q flow software (Medis medical imaging systems).

### Statistical analyses and graphical software

All data are reported as mean ± standard deviation of independent experimental replicates. Graphs and statistical analysis were made in GraphPad Prism. Surface response graphs and analysis were made in JMP (SAS, Cary, NC). Data normality was tested for all sample sets using Shapiro-Wilk test. P-values above 0.05 were considered non-significant. Sample sizes, statistical test used and p-values for each experiment are presented in the corresponding figure caption or in the corresponding supplementary table.

## AUTHOR CONTRIBUTIONS

J.A.B. Contributed to initial project idea, designed and conducted the experiments, analyzed the results, secured funding and wrote the manuscript. S.S. Modified and made devices for the pig experiments and contributed to 9-channel device perfusion experiments. H.E.O. Created the COMSOL models with their theoretical analysis and contributed to the 9-channel device perfusion experiments. D.V. imaged histology samples for the pig experiments. Y.R. Optimized and printed sugar constructs for the vascular network fabrication. H.W. Contributed to the endothelial and platelet adhesion assays and made the illustrations (Fig. 1a and 2a). K.C. contributed to an initial version of the COMSOL models. F.L. Imaged histology samples for the pig experiments. B.N.M. contributed to live/dead imaging of 9-channel devices. J.Z. and S.L. provided SC-islets for the pig experiments, edited the manuscript. T.J.K. Provided insight during the project, secured funding, edited the manuscript. J.I.T. Performed the pig surgery and helped design animal experiments. G.S. Performed doppler imaging and flow analysis in the porcine studies. R. L. L. Contributed to initial project idea, assisted with animal models and provided insight during the project, secured funding, edited the manuscript. A.B.D. contributed to initial idea and optimization of 9-channel sugar printing, provided insight during the project, secured funding, edited the manuscript. J.R. contributed to initial idea, designed the vascular bioreactor, contributed to the 9-channel sugar printing, provided insight during the project, secured funding. S.P. Performed the pig surgery, helped design animal studies, provided insight during the project and secured funding. C.A.H. Contributed to initial project idea, secured funding, provided insight during the project, analyzed results, wrote and edited the manuscript.

## ACKNOWLEDGMENTS

We thank Lisa Danielczak for supporting training and porcine studies, Emily Wilts for the transplantation of the cell-free material in mice, Robert K. Baker for providing the primers and methodology for qPCR and Kevin Bates for help with bi-axial tensile strength experiments. This study was supported by grants from the Canadian Institutes of Health Research (CIHR PJT-185939), JDRF (1-PNF-2016-249-S-B), Natural Sciences and Engineering Research Council of Canada (Discovery Grant 2018-06161), and the Quebec Network for Cell, Tissue and Gene Therapy – ThéCell (a thematic network supported by the Fonds de recherche du Québec–Santé (FRQS)). J.A.B. received a Vanier scholarship from CIHR and a doctoral scholarship from the Fonds de recherche du Québec nature et technologies (FRQNT). We also acknowledge the support provided by the following networks: the Cardiometabolic Health, Diabetes and Obesity – CMDO Research Network (thematic networks supported by the FRQS), PROTEO – The Quebec Network for Research on Protein Function, CQMF/QCAM -Quebec Centre for Advanced Materials (strategic teams supported by the FRQNT), the McGill Regenerative Medicine Network, the Montreal Diabetes Research Center, and the Canadian Biomaterials Society.

