## Supplementary Information for "Iterative sacrificial 3D printing and polymer casting to create complex vascular grafts and multi-compartment bioartificial organs"

### SUPPLEMENTARY FIGURES

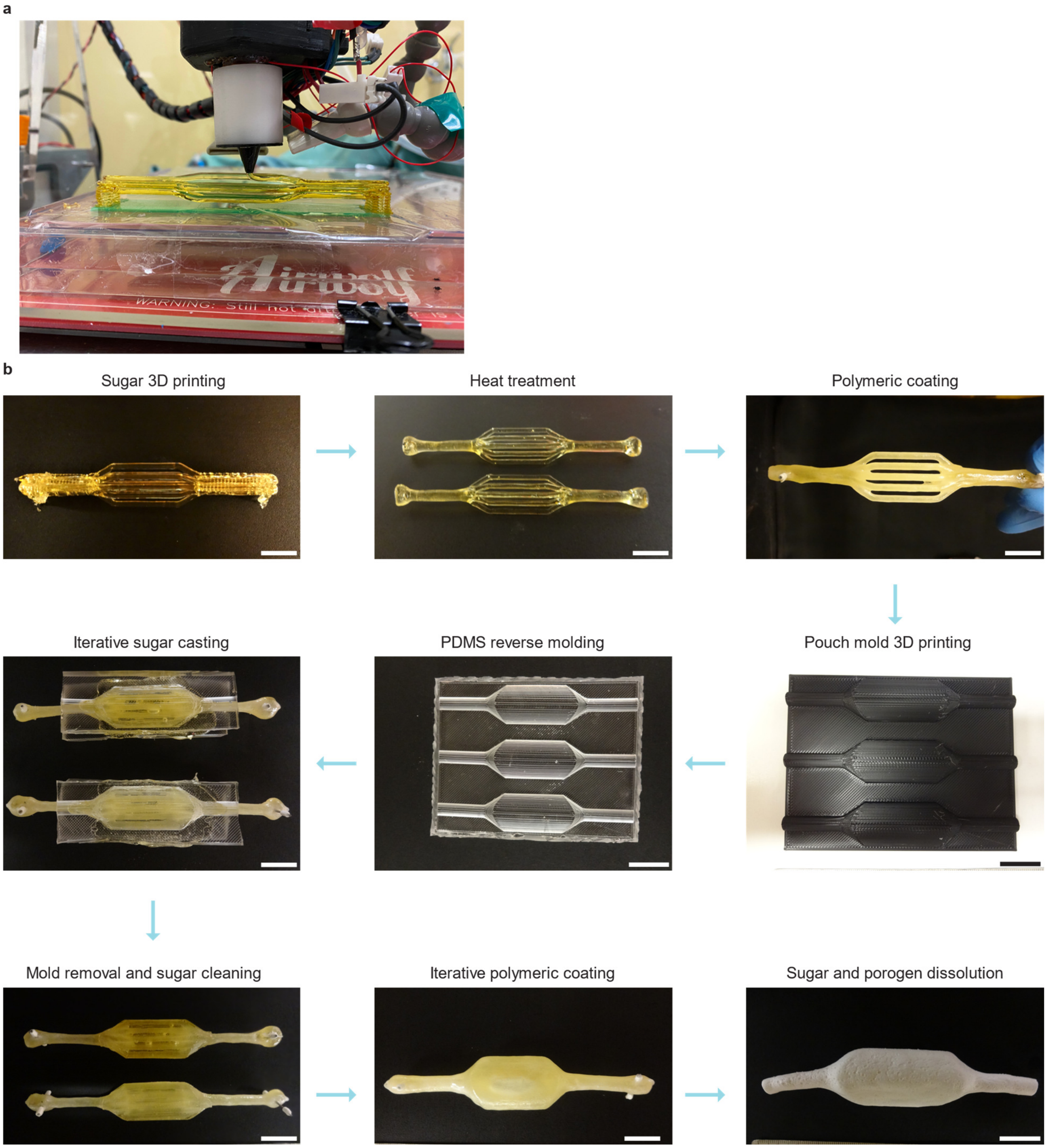


**Supplementary Figure 1 | Manufacturing methodology. a**, Representative image of the sugar printing process. **b**, Steps necessary for the fabrication of the vascular compartment (top). Steps for the fabrication of sugar molds for iterative molding and coating (middle). Steps for the fabrication of the external pouch of the device (bottom). Scale bars, 2 cm.

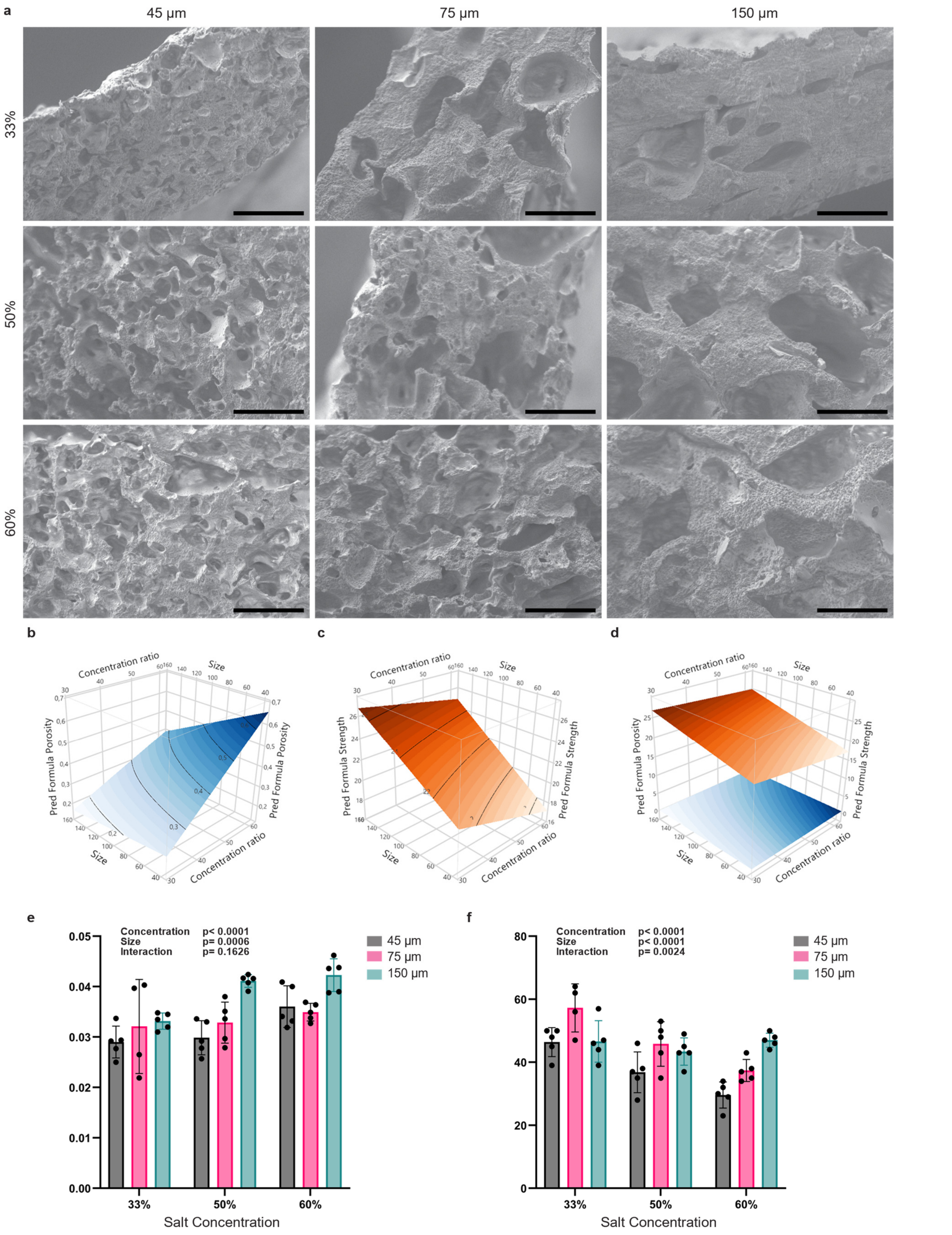
a
45 µm
75 µm
150 µm
33%
50%
60%
b
c
d
Concentration ratio
Size
Pred Formula Porosity
Pred Formula Strength
e
Concentration p< 0.0001
Size p= 0.0006
Interaction p= 0.1626
45 µm
75 µm
150 µm
33%
50%
60%
Salt Concentration
f
Concentration p< 0.0001
Size p< 0.0001
Interaction p= 0.0024
45 µm
75 µm
150 µm
33%
50%
60%
Salt Concentration

**Supplementary Figure 2 | Full factorial design.** **a**, Representative scanning electron microscopy images of the cross-section of the grafts in function of porogen size and concentration. Scale bars, 100 µm. **b**, Predicted surface response curve for the porosity of the grafts derived from a full factorial design with 5 measures for each point. **c**, Predicted surface response curve for the longitudinal tensile strength of the grafts derived from a full factorial design with 5 measures for each point. **d**, Graph allowing the visualization of the two surface response curves together. **e-f**, Graph showing polymer deposition (e) and diameter expansion when hydrated (f) for the grafts based on salt size and concentration. Mean and standard deviation of five different replicates is shown for each condition. Significance was determined using a two-way ANOVA with Tukey's multiple comparisons test. p-values for Tukey's test are in Supplementary Table 1.

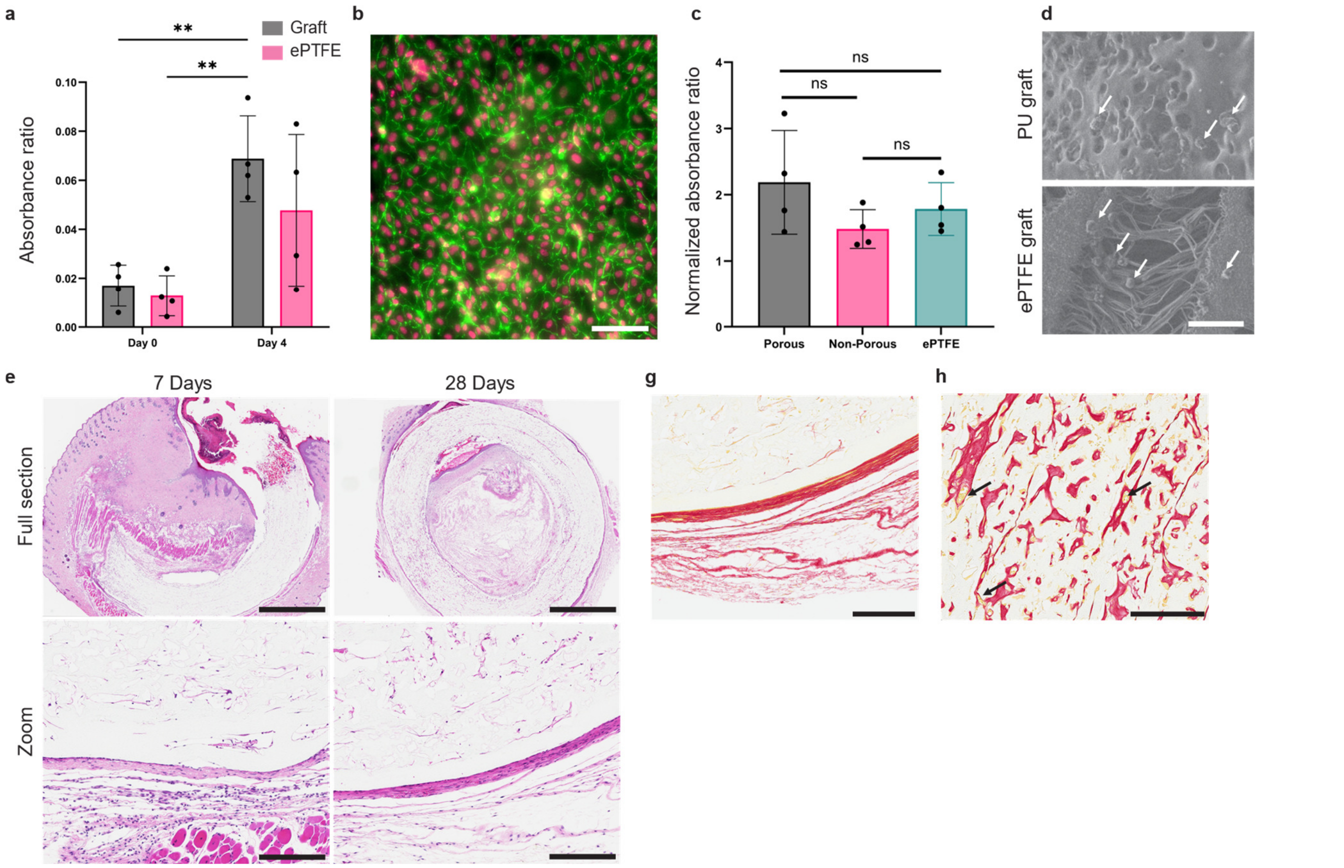


**Supplementary Figure 3 | Biological characterization of the grafts. a**, Graph of absorbance for WST-8 assay to assess the number of HUVECs adhering to the prosthesis and proliferating during static incubation. Mean and standard deviation of 4 independent biological replicates. **p < 0.01, determined by two-way ANOVA with Tukey's multiple comparisons test. **b**, Fluorescent images of confluent HUVECs on the internal surface of the prosthesis. Cells are labelled with DAPI (pink) and VE-cadherin (green). Scale bar, 50 µm. **c**, Graph of absorbance ratio for lactate dehydrogenase assay to assess the number of platelets adhering to the prosthesis during static incubation. Means and standard deviations of 4 different prostheses with the platelets of 4 different donors are shown. ns=non-significant, determined by one-way ANOVA with Tukey's multiple comparisons test. **d**, Electron micrograph of the PU and ePTFE graft surface after the platelet adhesion assay. White arrows show adhered platelets. Scale bar, 10 µm. **e**, Representative Hematoxylin and Eosin staining showing cross-sections of the grafts after subcutaneous transplantation in mice for 7 or 28 days. Scale bars, 2mm (top) and 200 µm (bottom). **f**, Representative Sirius Red staining showing cross-section of a graft after subcutaneous transplantation in mice for 28 days. Scale bar, 200 µm. **g**, Representative Sirius Red staining formation of putative vessels containing blood cells (yellow dots, black arrows) after subcutaneous transplantation in mice for 28 days. Scale bar, 200 µm.

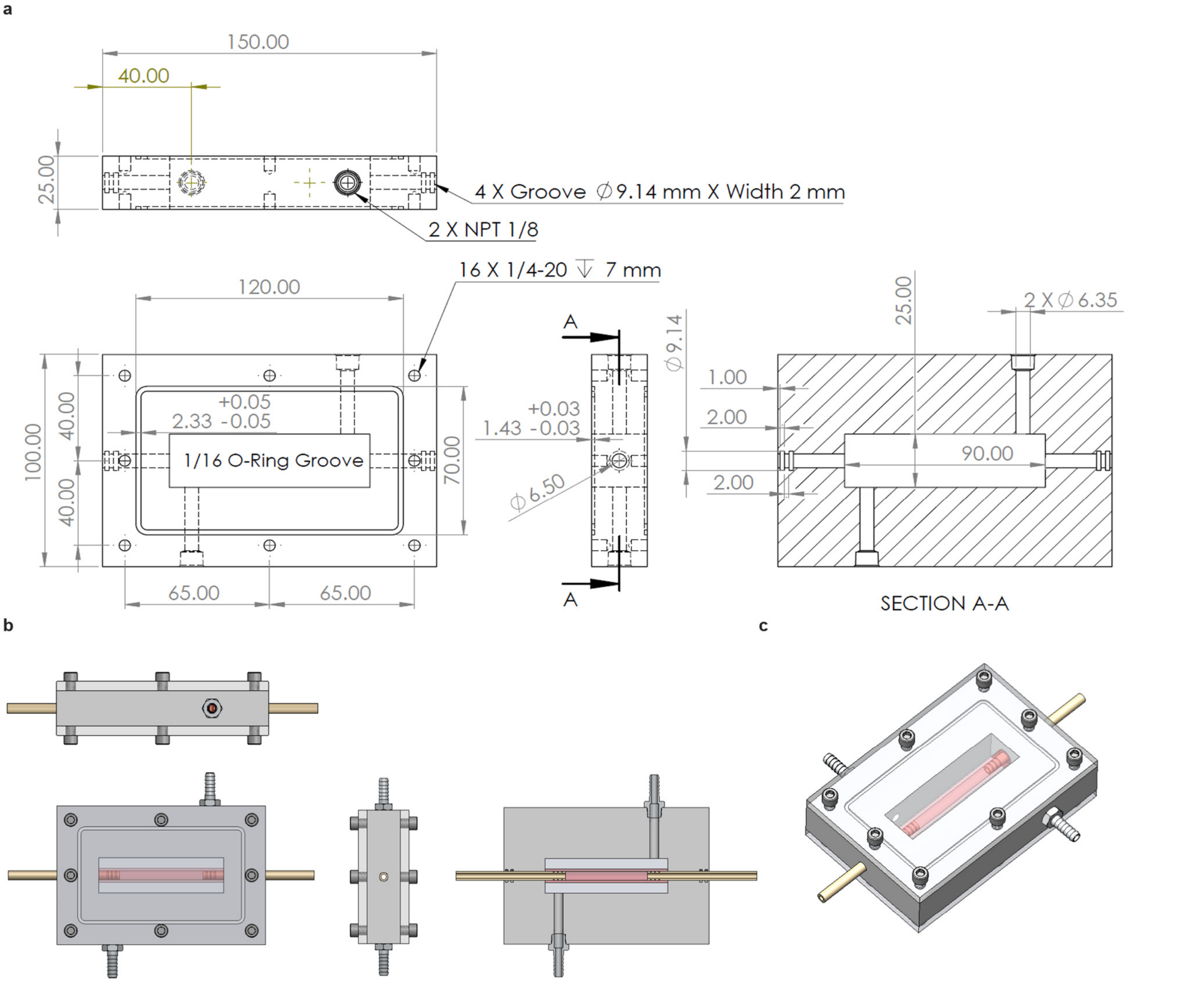


**Supplementary Figure 4 | Custom-made vascular chamber for perfusion. a**, Drawings of the vascular chamber with the different pieces needed and their dimensions. **b**, CAD illustration of the assembled bioreactor from different view. **c**, Illustration of the isometric view of the vascular chamber.

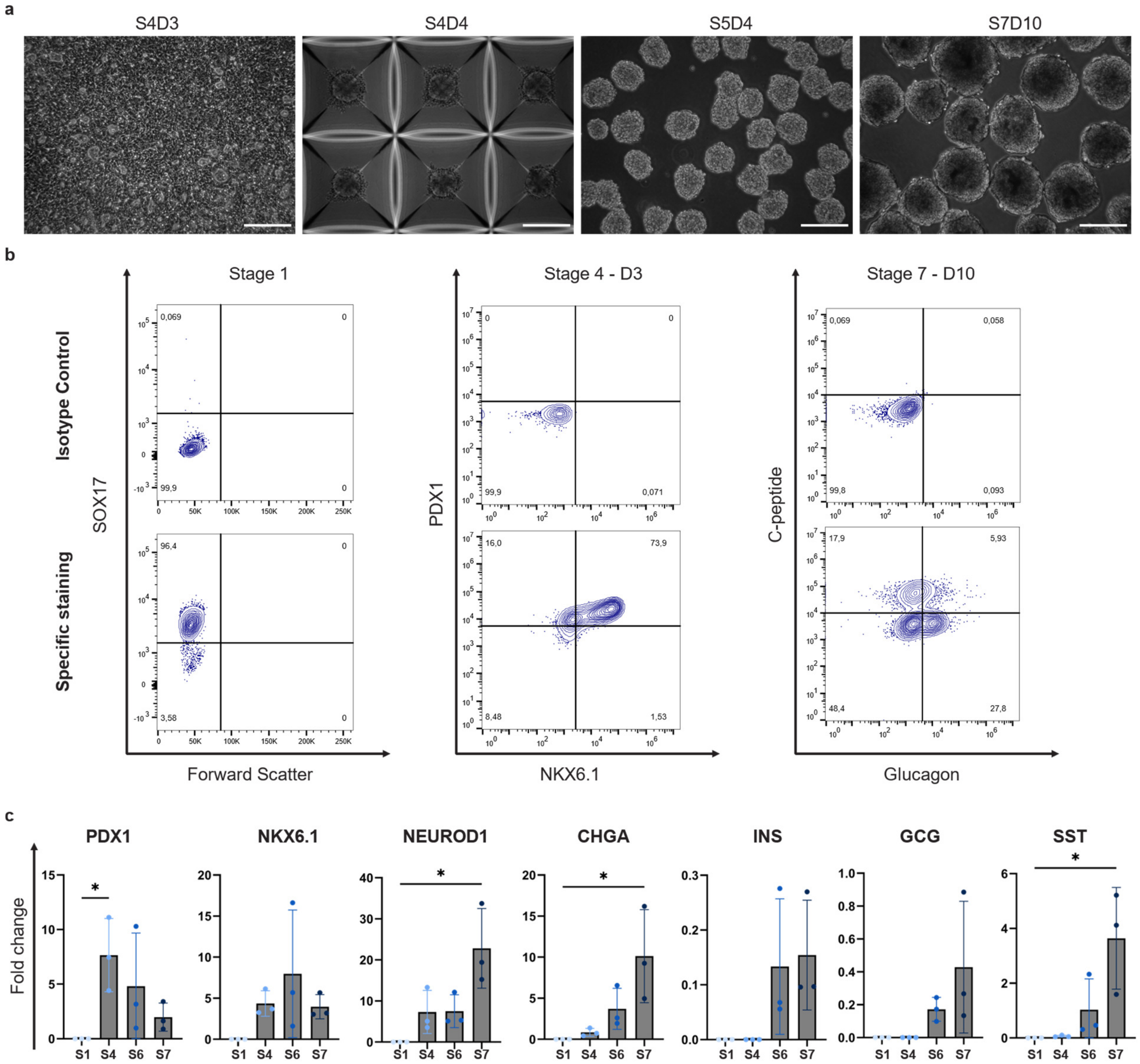


**Supplementary Figure 5 | SC-islet differentiation protocol characterization. a**, Representative brightfield images showing cell morphology at different stages of the differentiation protocol. Scale bars, 200 µm. **b**, Representative flow cytometry graphs showing population expression of genes of interest during the differentiation protocol. **c**, Gene expression profile of stem cell-derived cells during the differentiation protocol. For each gene at each stage, mean and standard deviation of 3 independent differentiation are shown (n=3 technical replicates for each point). *$p<0.05$, determined by one-way ANOVA with Dunn's multiple comparisons test. S1, Stage 1 day 3. S4, Stage 4 day 3. S6, Stage 6 day 7. S7, Stage 7 day 10.

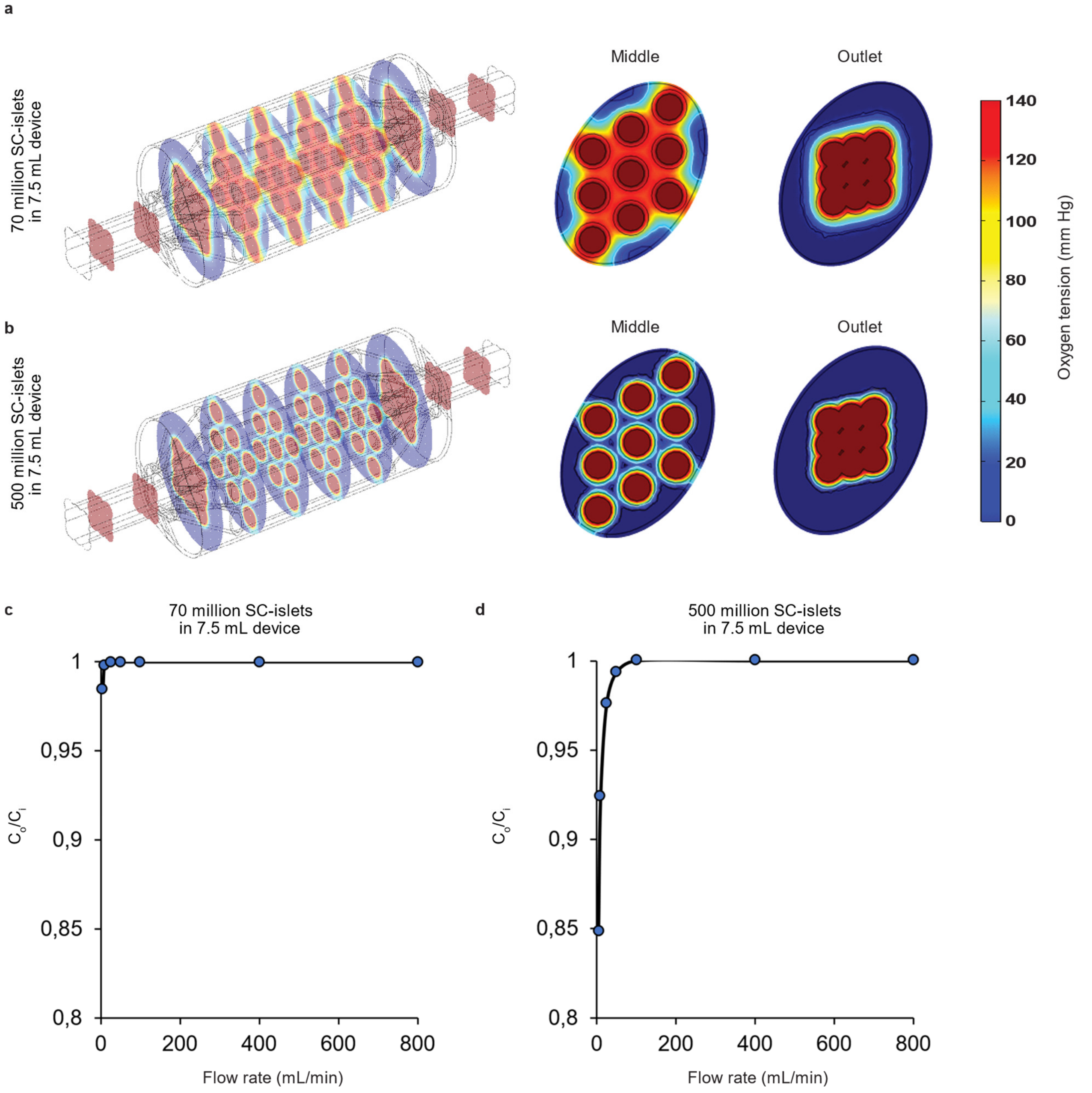


**Supplementary Figure 6 | Moving towards a vascularized bioartificial pancreas to accommodate a therapeutic dose of SC-islets**. **a-b**, Finite element model showing oxygen concentration profiles depending on cell density of SC-islets loaded within the device. Cross-section at the inlet and middle of the device are shown for the experimental dose used Fig. 5 (a, 70 million total) and therapeutic dose (b, 500 million cells per device). The white, grey and black contours represent the oxygen tension limits (20 mmHg, 10 mmHg and 0.1 mm Hg respectively). **c-d**, Effect of flow rate on the theoretical and experimental oxygen concentration at the outlet of the device at 70 million cells/device (c) or 500 million cells/device (d).

### SUPPLEMENTARY TABLES

*Table 1. Significance values for full factorial design*

| | Adjusted p values with significance symbol | | | | | | | |
|---|---|---|---|---|---|---|---|---|
| Tukey's multiple comparisons test | Polymer deposition | | Diameter expansion | | Strength | | Porosity | |
| 33%:45 vs. 33%:75 | ns | 0.9591 | ns | 0.0981 | ns | 0.4468 | ** | 0.0083 |
| 33%:45 vs. 33%:150 | ns | 0.7681 | ns | >0.9999 | ns | 0.4297 | ** | 0.0086 |
| 33%:45 vs. 50%:45 | ns | >0.9999 | ns | 0.1449 | ns | >0.9999 | **** | <0.0001 |
| 33%:45 vs. 50%:75 | ns | 0.831 | ns | >0.9999 | ns | >0.9999 | ns | 0.9345 |
| 33%:45 vs. 50%:150 | *** | 0.0008 | ns | 0.9926 | ns | 0.2204 | ns | 0.6735 |
| 33%:45 vs. 60%:45 | ns | 0.1537 | *** | 0.0006 | ns | 0.973 | **** | <0.0001 |
| 33%:45 vs. 60%:75 | ns | 0.3344 | ns | 0.2047 | ns | 0.986 | **** | <0.0001 |
| 33%:45 vs. 60%:150 | *** | 0.0002 | ns | >0.9999 | ns | 0.8917 | ns | 0.1336 |
| 33%:75 vs. 33%:150 | ns | >0.9999 | ns | 0.1106 | ns | >0.9999 | ns | >0.9999 |
| 33%:75 vs. 50%:45 | ns | 0.9946 | **** | <0.0001 | ns | 0.2459 | **** | <0.0001 |
| 33%:75 vs. 50%:75 | ns | >0.9999 | ns | 0.0676 | ns | 0.2045 | *** | 0.0003 |
| 33%:75 vs. 50%:150 | * | 0.0412 | * | 0.0129 | ns | >0.9999 | **** | <0.0001 |
| 33%:75 vs. 60%:45 | ns | 0.8612 | **** | <0.0001 | ns | 0.0557 | **** | <0.0001 |
| 33%:75 vs. 60%:75 | ns | 0.9755 | *** | 0.0001 | ns | 0.9528 | **** | <0.0001 |
| 33%:75 vs. 60%:150 | * | 0.0136 | ns | 0.1395 | ns | 0.997 | **** | <0.0001 |
| 33%:150 vs. 50%:45 | ns | 0.9206 | ns | 0.1284 | ns | 0.2338 | **** | <0.0001 |
| 33%:150 vs. 50%:75 | ns | >0.9999 | ns | >0.9999 | ns | 0.1939 | *** | 0.0002 |
| 33%:150 vs. 50%:150 | ns | 0.0686 | ns | 0.9887 | ns | >0.9999 | **** | <0.0001 |
| 33%:150 vs. 60%:45 | ns | 0.9639 | *** | 0.0005 | ns | 0.0522 | **** | <0.0001 |
| 33%:150 vs. 60%:75 | ns | 0.9983 | ns | 0.183 | ns | 0.9463 | **** | <0.0001 |
| 33%:150 vs. 60%:150 | * | 0.0224 | ns | >0.9999 | ns | 0.9962 | **** | <0.0001 |
| 50%:45 vs. 50%:75 | ns | 0.9526 | ns | 0.2047 | ns | >0.9999 | *** | 0.0001 |
| 50%:45 vs. 50%:150 | ** | 0.0022 | ns | 0.5921 | ns | 0.1033 | *** | 0.0006 |
| 50%:45 vs. 60%:45 | ns | 0.2897 | ns | 0.4789 | ns | 0.9984 | *** | 0.0002 |
| 50%:45 vs. 60%:75 | ns | 0.5397 | ns | >0.9999 | ns | 0.9085 | ns | 0.2902 |
| 50%:45 vs. 60%:150 | *** | 0.0006 | ns | 0.1 | ns | 0.6986 | * | 0.0118 |
| 50%:75 vs. 50%:150 | ns | 0.0519 | ns | 0.9984 | ns | 0.0828 | ns | 0.9997 |
| 50%:75 vs. 60%:45 | ns | 0.9369 | *** | 0.001 | ns | 0.9995 | **** | <0.0001 |
| 50%:75 vs. 60%:75 | ns | 0.9951 | ns | 0.2808 | ns | 0.87 | **** | <0.0001 |
| 50%:75 vs. 60%:150 | * | 0.0164 | ns | >0.9999 | ns | 0.6353 | ns | 0.8005 |
| 50%:150 vs. 60%:45 | ns | 0.5345 | ** | 0.0072 | * | 0.0188 | **** | <0.0001 |
| 50%:150 vs. 60%:75 | ns | 0.2858 | ns | 0.7041 | ns | 0.7827 | **** | <0.0001 |
| 50%:150 vs. 60%:150 | ns | >0.9999 | ns | 0.9765 | ns | 0.9498 | ns | 0.9777 |
| 60%:45 vs. 60%:75 | ns | >0.9999 | ns | 0.3731 | ns | 0.5157 | ns | 0.116 |
| 60%:45 vs. 60%:150 | ns | 0.2707 | *** | 0.0004 | ns | 0.2746 | **** | <0.0001 |
| 60%:75 vs. 60%:150 | ns | 0.1182 | ns | 0.1449 | ns | >0.9999 | **** | <0.0001 |

*Table 2. Differentiation medium recipes*

<table>
<tr><th>Medium</th><th>Basal Medium</th><th>Soluble Factors</th></tr>
<tr><td>S1</td><td rowspan="2">MCDB131 + 10mM glucose + 1.5 g/L $NaHCO_3$ + 0.5% fatty acid free bovine serum albumin (FAF-BSA) + 1x GlutaMAX + 1% Pen/Strep</td><td>+ 100 ng/mL Activin A<br>+ 3 µM CHIR99021 (Day1 only)</td></tr>
<tr><td>S2</td><td>+ 0.25 mM ascorbic acid<br>+ 50 ng/mL keratinocyte growth factor (KGF)<br>+ 1.25 µM IWP-2</td></tr>
<tr><td>S3</td><td rowspan="2">MCDB131 + 10mM glucose + 2.5 g/L $NaHCO_3$ + 2% FAF-BSA + 1X GlutaMAX + 1% Pen/Strep</td><td>+ 0.25 mM ascorbic acid<br>+ 1:200 insulin-transferrin-selenium-ethanolamine (ITS-X)<br>+ 50 ng/mL KGF<br>+ 0.25 µM SANT-1<br>+ 1 µM retinoic acid<br>+ 100 nM LDN193189<br>+ 200 nM TPB (PKC activator)</td></tr>
<tr><td>S4</td><td>+ 0.25 mM ascorbic acid<br>+ 1:200 ITS-X<br>+ 2 ng/mL KGF<br>+ 0.25 µM SANT-1<br>+ 0.1 µM retinoic acid<br>+ 200 nM LDN193189<br>+ 100 nM TPB</td></tr>
<tr><td>S5</td><td>MCDB131 + 20mM glucose + 1.5 g/L $NaHCO_3$ + 2% FAF-BSA + 1x GlutaMAX + 1% Pen/Strep</td><td>+ 1:200 ITS-X<br>+ 10 µM zinc sulfate<br>+ 10 µg/mL heparin<br>+ 0.25 µM SANT-1<br>+ 1 µM T3<br>+ 100 nM Gamma secretase inhibitor XX<br>+ 10 µM ALK5 inhibitor II<br>+ 0.05 µM retinoic acid<br>+ 100 nM LDN193189</td></tr>
<tr><td>S6</td><td>MCDB131 + 20mM glucose + 1.5 g/L $NaHCO_3$ + 2% FAF-BSA + 1x GlutaMAX + 1% Pen/Strep</td><td>+ 1:200 ITS-X<br>+ 10 µM zinc sulfate<br>+ 10 µg/mL heparin<br>+ 1 µM T3<br>+ 100 nM Gamma secretase inhibitor XX<br>+ 10 µM ALK5 inhibitor II<br>+ 100 nM LDN193189</td></tr>
</table>

| | | |
|---|---|---|
| S7 | MCDB131 + 20mM glucose + 1 g/L $NaHCO_3$ + 2% FAF-BSA + 1x GlutaMAX + 1% Pen/Strep + 1x non-essential amino acids +1x Trace element A + 1x Trace element B | + 1:200 ITS-X<br>+ 10 µM zinc sulfate<br>+ 10 µg/mL heparin<br>+ 1 µM T3<br>+ 1mM N-acetyl cysteine |

*Table 3. List of primers*

| Name | Sequence |
|---|---|
| hCHGA-F | CCTGTGAACAGCCCTATGAATAAAG |
| hCHGA-R | CTGATGTCTCAGAATGGAAAGGATC |
| hGCG-F | TTCTACAGCACACTACCAGAAGA |
| hGCG-R | CTGGGAAGCTGAGAATGATCTG |
| hINS-F | GCAGCCTTTGTGAACCAACA |
| hINS-R | GGTGTGTAGAAGAAGCCTCGTT |
| hNeuroD1-F | GGTTATGAGACTATCACTGCTCAG |
| hNeuroD1-R | AGAACTGAGACACTCGTCTGTC |
| hNKX6.1-F | CCTGTACCCCTCATCAAGGAT |
| hNKX6.1-R | CAAGTATTTTGTTTGTTCGAAAGTCTTCT |
| hPdx1-F | CCCTCTTTTAGTGATACTGGATTGG |
| hPdx1-R | CCTTCCAATGTGTATGGTACAGTTTC |
| hSST-F | TCCGTCAGTTTCTGCAGAAGTC |
| hSST-R | CTGGGACAGATCTTCAGGTTCC |
| hNFX1-F | TTTCAGAACAAAGGAGCTTCCAT |
| hNFX1-R | CTTATCCACACAGCATATCTCATTACA |

Table 4. Oxygen transport modelling parameters

| Parameter | Value | Reference |
|---|---|---|
| MIN6 Maximum oxygen consumption rate (OCR, $R_{O_2,M6}$) | 0.129 mol $s^{-1}$ $m^{-3}$ | 60 |
| MIN6 Monod constant for oxygen consumption ($K_{s,M6}$) | 0.62 µM | 61,62 |
| Necrotic oxygen tension ($C_{cr,M6}$) | 0.1 mmHg | 61 |
| Islet maximum oxygen consumption rate (OCR, $R_{O_2,HI}$) | 0.034 mol $s^{-1}$ $m^{-3}$ | 63 |
| Islet Monod constant for oxygen consumption ($K_{s,HI}$) | 1 µM | 63 |
| Oxygen diffusivity in alginate ($D_{alg}$) | $2.5 \times 10^{-5}$ $cm^2$ $s^{-1}$ | 64 |
| Oxygen diffusivity in cells ($D_{cells}$) | $1.24 \times 10^{-5}$ $cm^2$ $s^{-1}$ | 61 |
